# Zebrafish screen of schizophrenia risk genes reveals convergent dysregulation of cholesterol metabolism

**DOI:** 10.64898/2026.02.17.706411

**Authors:** Anna J. Moyer, Mary E.S. Capps, Claire L. Conklin, Brandon L. Bastien, Verdion Martina, William C. Gannaway, Camden E. Cummings, Jaqueline A. Martinez, Mandy Chen, Gretchen Kioschos, Emma G. Torija-Olson, Morgan C. Klein, Michael D. Vivian, Caleb C.S. Calhoun, Summer B. Thyme

## Abstract

Rare coding variants provide a tractable entry point for understanding the molecular mechanisms underlying schizophrenia risk. Here, we generated and characterized zebrafish lines with mutations in the orthologs of >20 human schizophrenia-associated genes, including eight of the top ten SCHEMA genes, genes disrupted in childhood-onset schizophrenia (COS), and genes located within recurrent copy number variants. Whole-brain phospho-Erk activity mapping and behavioral profiling identified phenotypes in multiple mutant lines. We prioritized a protein-truncating mutation in *sp4*, which encodes an activity-dependent transcription factor, and a COS-associated missense mutation in *atp1a3a*, which encodes a Na+/K+ ATPase pump, for additional characterization. Both knockout and point mutations in *atp1a3a* disrupted brain activity and behavior in larvae and impaired navigation of a Y-maze in juveniles. Bulk RNA sequencing data from adult *sp4* and *atp1a3a* brains highlighted convergent upregulation of sterol biosynthesis pathways, including increased expression of *srebf2* and *msmo1.* Analysis of previously published telencephalon single-cell data demonstrated that cholesterol synthesis genes are enriched in astrocyte-like cells and increase in expression during post-larval development. Consistent with transcriptomic findings, filipin staining indicated increased free cholesterol in juvenile *sp4* and *atp1a3a* mutant brains. Our findings identify dysregulation of glial and sterol-associated programs as a shared molecular consequence of two distinct schizophrenia risk mutations. Although whether sterol pathway dysregulation represents a primary pathogenic mechanism or a secondary response to changes in neuronal activity requires further investigation, the convergence observed between genetic models and developmental stages suggests that disruptions to lipid homeostasis could represent a shared feature of schizophrenia disease biology.

## Introduction

Schizophrenia is a highly heritable psychiatric disorder, with genetics estimated to account for as much as 60-80% of its risk^1,2^. The search for the underlying genetic causes began several decades ago, with notable examples such as a rare, strong-effect chromosomal translocation disrupting *DISC1*^3,4^, implicating variation in this gene as conferring schizophrenia susceptibility. Genetic mapping of schizophrenia accelerated dramatically following the advent of genome-wide association studies (GWAS) from large international consortia, resulting in the identification of nearly 300 genome-wide significant loci thus far^5^. A main conclusion of these studies is that schizophrenia is highly polygenic, with risk distributed across thousands of common variants that individually contribute small effects. The vast majority of variants lie in noncoding, extragenic regions and presumably act through transcriptional regulation^6^. Consequently, only a small fraction of loci have been confidently linked to target genes, making signals difficult to interpret mechanistically or translate into animal models.

Because schizophrenia GWAS signals involve human regulatory variation with small individual effect sizes, functional follow-up in non-human models is challenging. Human genomic studies have revealed some convergent signals, including those related to synaptic pruning^7,8^ and excitatory signaling (e.g., NMDA receptors and voltage-gated calcium channels), but mechanistic connections are largely limited to well-characterized genes and pathways. To define molecular pathways underlying schizophrenia, case-control iPSC and brain organoid models preserve this genetic variation^9,10^. Such studies remain difficult, however, because large samples are necessary to gain adequate insight given heterogeneous genetic backgrounds, while cost limits scalability^11^. Further, even the most advanced culture systems only represent a limited subset of brain cell types and lack mature, organized circuits. Large-scale postmortem cortical transcriptomic efforts have uncovered reproducible dysregulation of gene programs associated with synaptic development and neuronal activity^12^, as well as cell type-specific effects^13,14^. Postmortem tissue, however, inherently provides only an end-stage snapshot and cannot resolve developmental perturbations. Neither human-specific approach accurately recapitulates circuit-level dysfunction or non-autonomous cellular interactions. Animal models therefore remain essential, but their generation is more compatible with non-GWAS forms of genetic risk.

Zebrafish offer a tractable animal model for defining the neurodevelopmental consequences of many genetic perturbations. Zebrafish share approximately 70% of their genes with humans^15^, have similar neuron types^16,17^, and fit into individual wells of 96-well plates at the larval stage. Their small size makes them ideally suited for drug screening; multiple compounds discovered using zebrafish are in clinical trials^18,19^, including for Dravet Syndrome^20^, a severe genetic form of epilepsy. To investigate the pathways underlying schizophrenia, we previously generated presumed loss-of-function, protein-truncating mutants for zebrafish orthologs of 132 human genes located within or near GWAS loci^21^. The study uncovered numerous mutants with relevant behavioral and brain activity phenotypes, such as decreased prepulse inhibition, which nominated likely candidate genes in multi-gene loci. A similar zebrafish survey of 22 of the 45 genes in the 22q11.2 locus, deletion of which is one of the strongest genetic risk factors for schizophrenia, highlighted mitochondrial function in neural stem cells as a shared phenotype^22^. A key advantage to the zebrafish-based approach is the ability of a single laboratory to systematically compare many genetic perturbations using standardized assays and environmental conditions. Despite these advantages, the lack of face validity of these knockouts limits their use to primarily hypothesis generation and basic investigation rather than drug screening to mitigate phenotypes.

Here, we sought to model additional sources of genetic variation that contribute to schizophrenia risk with larger effect sizes than common variants identified by GWAS. Rare coding mutations identified through exome sequencing provide tractable entry points for developing *in vivo* models that more closely approximate molecular aspects of human disease. We therefore generated and characterized zebrafish lines representing more than 20 human genes, including eight of the top ten SCHEMA^23^ hits, ultrarare alleles identified in childhood-onset schizophrenia (COS) cases, and genes within small copy number variants (CNVs). Based on larval brain activity, brain structure, and behavioral screening, we prioritized two lines for in-depth analysis: a protein-truncating allele of *sp4* and a missense substitution in *atp1a3a*, orthologous to the V129M mutation identified in an individual with COS^24^. Adult brain transcriptomics revealed shared dysregulation of cholesterol metabolism, marked by altered expression of astrocyte-enriched genes associated with the developmental transition in brain cholesterol synthesis, and corroborated by juvenile brain imaging. As neither gene has previously been implicated in this pathway, these findings suggest that diverse, large-effect schizophrenia risk mutations can converge on overlapping molecular mechanisms.

## Results

### Selection and generation of zebrafish mutants for schizophrenia risk genes

We selected genes for zebrafish mutagenesis from multiple non-GWAS sources, and all mutants were subjected to our previously established phospho-Erk (pErk)-based whole-brain activity and structure mapping and a behavioral pipeline^21,25^ (**Supplementary Table S1**). We included eight genes from SCHEMA, a large exome-sequencing consortium that identified ten genes surpassing significance for rare coding variant burden in schizophrenia^23^. Mutations in these genes confer an approximately 3- to 50-fold increase in schizophrenia risk, making them strong candidates for animal modeling. We also included genes implicated in childhood-onset schizophrenia (COS), which are associated with severe early-onset disease and include both point mutations and genes disrupted by small copy number variants^24,26–28^. Although we mainly generated typical protein-truncating alleles, we did include specific amino acid substitutions for some COS variants, in particular, the *ATP1A3* V129M allele^24^. To model 15q11.2 BP1–BP2 microdeletion syndrome^29,30^, we removed a >130 kilobase region encompassing orthologs of the four recurrently deleted genes (*TUBGCP5*, *CYFIP1*, *NIPA1*, *NIPA2*), which are syntenic in zebrafish. Loss of either *nipa1* or *nipa2* individually resulted in reduced brain activity (**Supplementary Fig. S1**). However, homozygous analysis of the full deletion was precluded by severe morphological defects caused by homozygous loss of *cyfip1*. Additional mutants for several genes from other rare, small CNVs^28,31,32^ (*PDXDC1*, *NTAN1*, *CHRNA7*, *OTUD7A*) or individual case studies (*PTPRG*^33^, *FSTL5*^34,35^), failed to produce detectable alterations in brain activity (**Supplementary Fig. S1**, **Supplementary Fig. S2**) or behavior (**Supplementary Fig. S3**, **Supplementary Fig. S4**). Lastly, in an attempt to more closely approximate GWAS loci, we removed intronic genomic regions that align with the areas implicated by GWAS, as these lines would have better construct validity than protein-truncation mutations^21^. This effort also did not yield phenotypes (**Supplementary Fig. S5**). From this diverse set of alleles, SCHEMA and COS genes produced the strongest effects, described below.

### Phenotypic analysis of SCHEMA schizophrenia risk genes

The top ten SCHEMA genes represent the first set of exome mutations that have reached population-wide significance and confer a magnitude of risk comparable to recurrent CNVs such as 22q11.2 deletions. Of these ten genes, we excluded *GRIN2A* and *GRIA3*; we had already generated and characterized the *grin2aa/b* mutants^21^, as this gene is in a consistently associated GWAS locus^36^. Although we had not previously studied *GRIA3*, *gria1a/b* mutants were in our published mutant set, and we chose to focus on the eight more functionally diverse genes. We were unable to generate a true protein-truncating allele for *setd1a* and we instead analyzed an approximately 200 amino acid in-frame deletion of a conserved protein region.

Whole-brain activity and structure maps were compared to brain regions from the Z-Brain atlas^37^ (**Fig. 1a**). Knockout of both zebrafish orthologs of *CUL1* and *TRIO* resulted in embryonic lethality; therefore, comparisons could not include the double homozygous and, given overlapping roles of the ohnologs and genetic compensation^38^, are likely incomplete knockouts. The strongest phenotypes were observed for *setd1a* and *rb1cc1* mutants. Increased activity was dominant throughout the brain (**Fig. 1b**, **Supplementary Fig. S6**), and structural analysis, calculated using deformation-based morphometry^21,39^, revealed substantial brain enlargement in *setd1a* and *rb1cc1* mutants (**Fig. 1c**, **Supplementary Fig. S7**). Among the remaining lines, *cacna1g* and *xpo7* exhibited no detectable phenotypes. The *herc1* mutants displayed mildly reduced brain activity, particularly in the cerebellum and hindbrain (**Fig. 1d**). Brain activity differences in the *sp4* mutants were localized to the telencephalon, similar to several mutants in our previous study^21^.

**Fig. 1:**
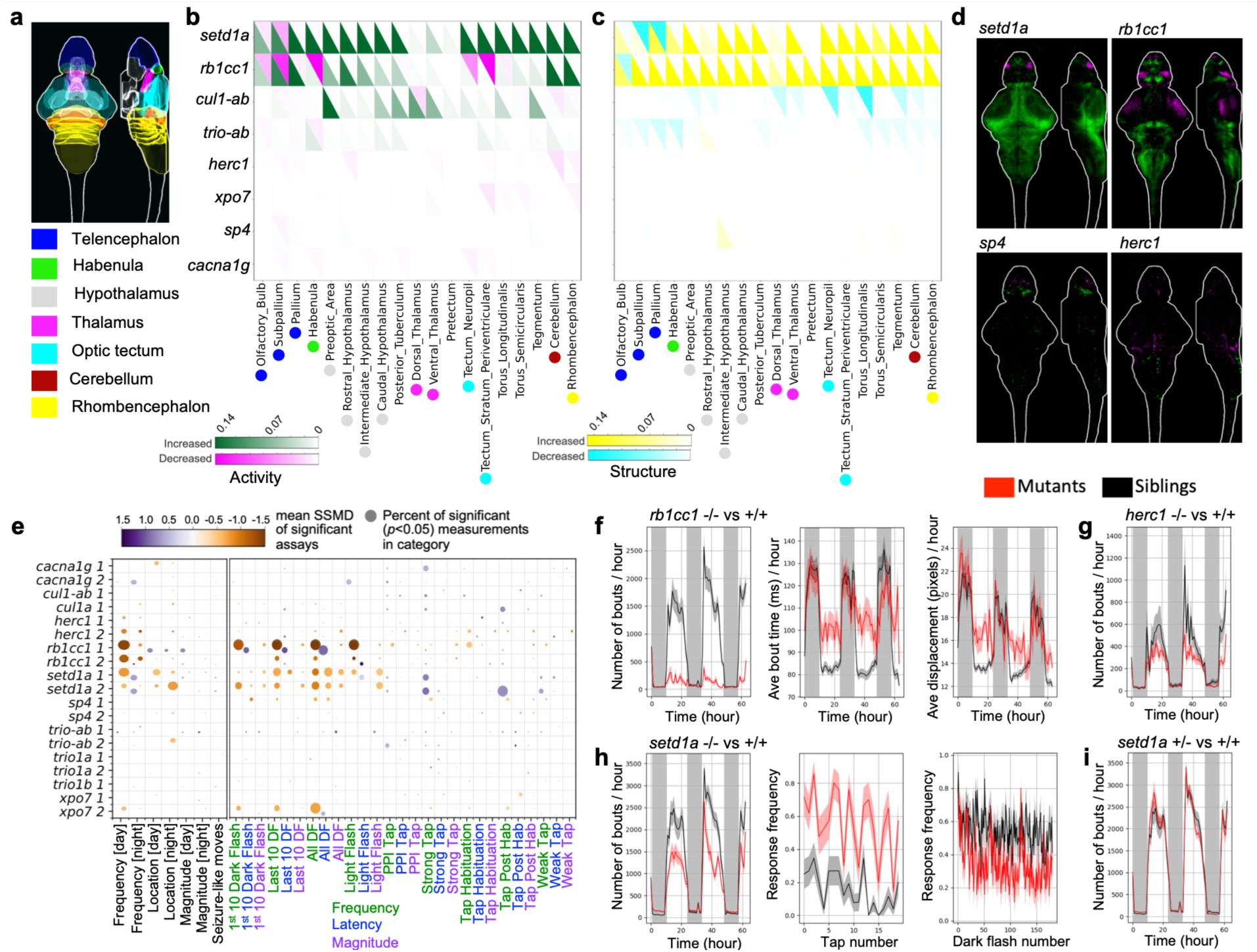
Whole-brain activity, morphology, and behavior phenotypes in SCHEMA gene mutants. **a**) Location of selected major brain regions in the zebrafish brain, based on the Z-Brain atlas. **b**) Summary of pErk comparisons between groups, where magenta represents decreased activity and green represents increased. The signal in each region was summed and divided by the total size of that region. Displayed data are the following comparisons from the run1 set: homozygous−/− versus heterozygous +/− and wildtype +/+ siblings for single gene mutants, −/−;+/− versus +/+;+/− for *cul1-ab*, and −/−;+/−versus +/+;+/− for *trio-ab*. The activity brain maps of each genotype comparison are available in Supplementary Fig. S6. **c**) Summary of structure comparisons between groups, where cyan represents decreased size and yellow represents increased. The structure brain maps of each genotype comparison are available in Supplementary Fig. S7. **d**) Examples of activity differences in mutants displayed as sum-of-slices projections (Z- and X-axes). Brain images represent the significant differences in signal between the homozygous −/− and wildtype sibling +/+ groups. **e**) Visualization summary of behavioral phenotypes. The size of the bubble represents the percent of significant measurements in the summarized category, and the color represents the mean of the strictly standardized mean difference (SSMD) of the significant assays in that category. Biological replicates from independent embryo batches are shown, comparing homozygous −/− to wildtype +/+ siblings except for *cul1-ab* (−/−;+/− versus +/−;+/−) and *trio-ab* (+/−;−/− versus +/−;+/−). The N for all experiments is available in Table S1. The comparisons of each genotype separately are available in Supplementary Fig. S8 and Supplementary Fig. S9. Baseline behavior: frequency of movement (e.g., number of bouts / hour), magnitude of movement (e.g., bout distance traveled), and location (e.g., fraction of bout in center of well) preferences were calculated for the baseline data. Stimulus-driven behavior: response frequency, latency, and magnitude for dark flashes (DF), light flashes, acoustic habituation (Hab), and acoustic stimulation that is strong, weak, or strong preceded by weak (PPI). **f**) Examples of *rb1cc1* baseline behavioral phenotypes. *p* values: 9.5e-8 for bout frequency (number of bouts / hour), 0.0622 for bout time for the entire time course and 0.0043 for the day 1 data subsection (not shown: day1ppihab, binned by 10-minutes), and 0.034 for bout displacement. **g**) Example of *herc1* baseline frequency of movement phenotype. *p* value: 0.017. **h**) Examples of *setd1a* behavioral phenotypes. *p* values: 0.00014 for bout frequency (number of bouts / hour), 2.4e-6 for strong tap response frequency at night at 6 dpf (sound frequency 1400 Hz), 0.0001 dark flash response frequency at 6 dpf (all dark flashes). **i**) No strong phenotype in the *setd1a* +/− versus +/+ groups, except a weak increased movement frequency on the day 0 night subsection (*p* value: 0.0053, binned per minute). Replicate data for f-i are available in Supplementary Fig. S10. Plots in f-i represent mean ± s.e.m. *p* values are Kruskal-Wallis ANOVA. The N for all experiments is available in Supplementary Table S1.

Behavioral outcomes largely aligned with the extent of phenotypes observed in brain imaging. As expected based on their dramatic brain size and activity differences, the *rb1cc1* and *setd1a* mutants exhibited far stronger behavioral alterations than any other lines (**Fig. 1e**, **Supplementary Fig. S8**). The number of movement bouts was far lower for *rb1cc1* mutants than sibling controls, and the magnitude of these rarer movements was increased (e.g., time and displacement) (**Fig. 1f**). The *herc1* mutants also consistently displayed fewer movement bouts (**Fig. 1g**), in line with the reduced activity in the hindbrain (**Fig. 1d**), although to a much lesser extent than *rb1cc1* mutants. Daytime locomotor activity was reduced in *setd1a* mutants, but nighttime movement frequency was increased relative to sibling controls (**Fig. 1h**). Both *rb1cc1* and *setd1a* mutants responded less often to dark flashes, but *setd1a* mutants responded more often to acoustic stimulation (**Fig. 1e**, **Fig. 1h**, **Supplementary Fig. S9**), suggesting a more complex change in circuitry that does not simply overall dampen activity and responsiveness. Overall, *setd1a* heterozygotes exhibited minimal phenotypes (**Fig. 1i**), with the exception of a slight increase in night movement frequency, indicating that in-frame removal of the protein domain did not yield a dominant negative outcome.

### Phenotypic analysis of rare childhood-onset schizophrenia (COS) risk genes

Exome sequencing of trios has revealed a small number of missense *de novo* variants potentially contributing to COS. Although this disorder is exceedingly rare, four individuals have been identified with distinct variants in *ATP1A3*^24,27^. From a cohort of 17 cases, several genes linked to neuronal function were nominated^26^. This variant set included *PPP1R13B*, which is located in an associated locus from GWAS^36^. From these previously identified genes, we selected several to make protein-truncating mutations as well as three specific amino acid substitutions (**Supplementary Table S1**).

Whole-brain activity and structure maps were compared to brain regions from the Z-Brain atlas^37^ (**Fig. 2a**). In spite of the human genetic links for these alleles, the only mutants with robust phenotypes were *atp1a3a* and *atp1a3b*, orthologs of *ATP1A3* (**Fig. 2b**, **Fig. 2c**, **Supplementary Fig. S11**, **Supplementary Fig. S12**). Both the *atp1a3a* missense mutation, corresponding to the human V129M variant, and protein-truncating lines exhibited reduced brain activity throughout the brain (**Fig. 2d**). In addition, the protein-truncating mutation (*atp1a3a-ko*) had strongly increased activity in the lateral telencephalon and reduced brain size (**Fig. 2c**, **Fig. 2d**). The knockout allele of the *atp1a3b* ortholog also affected brain activity and structure, although much less so than for *atp1a3a*.

**Fig. 2:**
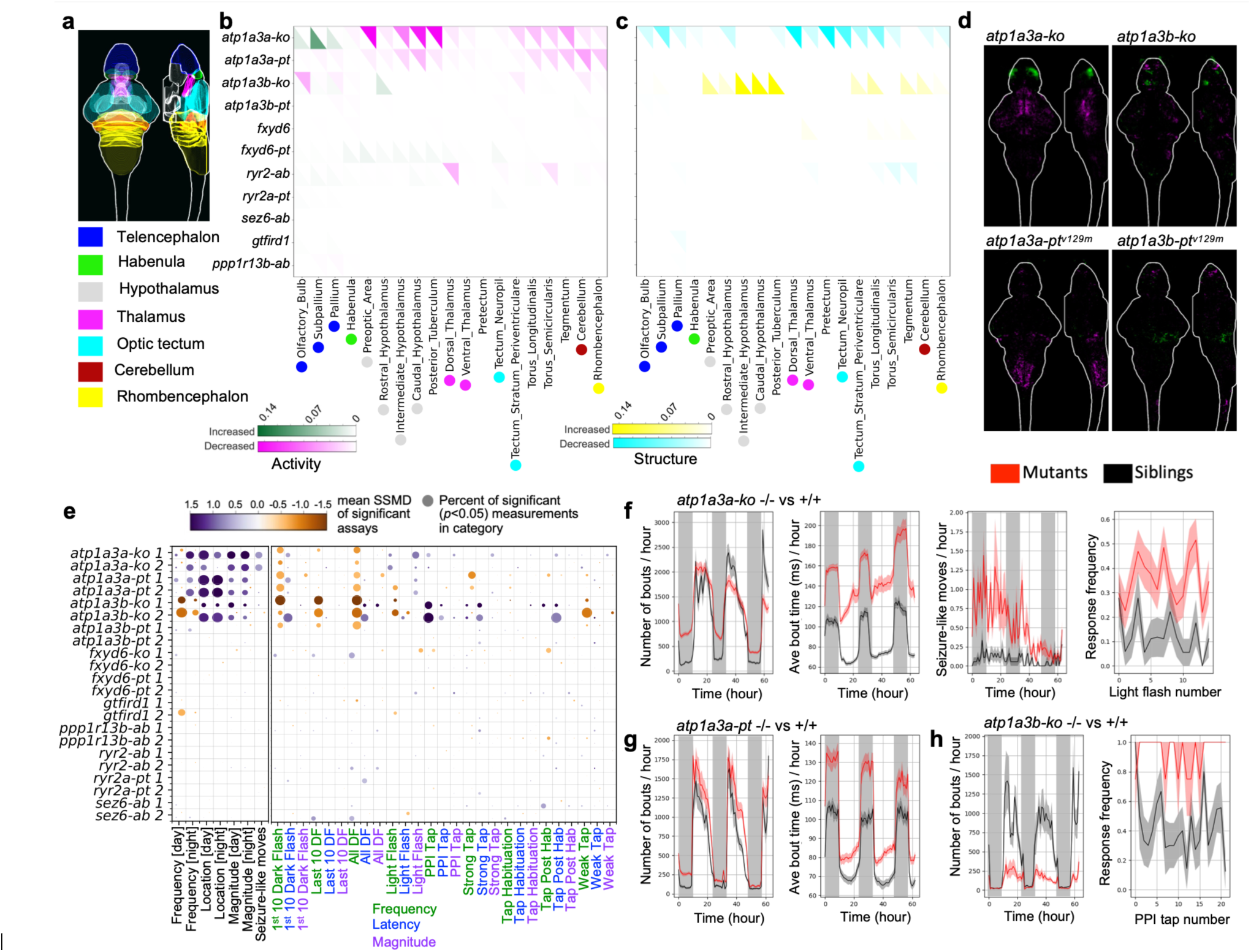
Whole-brain activity, morphology, and behavior phenotypes in COS gene mutants. **a**) Location of selected major brain regions in the zebrafish brain, based on the Z-Brain atlas. **b**) Summary of pErk comparisons between groups, where magenta represents decreased activity and green represents increased. The signal in each region was summed and divided by the total size of that region. Displayed data are the following comparisons from the run1 set: homozygous −/−versus heterozygous +/− and wildtype +/+ siblings for single gene mutants, −/−;−/− versus +/−;+/− for duplicated a/b genes. The activity brain maps of each genotype comparison are available in Supplementary Fig. S11. **c**) Summary of structure comparisons between groups, where cyan represents decreased size and yellow represents increased. The structure brain maps of each genotype comparison are available in Supplementary Fig. S12. **d**) Examples of activity differences in mutants displayed as sum-of-slices projections (Z- and X-axes). Brain images represent the significant differences in signal between the homozygous −/− and wildtype sibling +/+ groups. **e**) Visualization summary of behavioral phenotypes. The size of the bubble represents the percent of significant measurements in the summarized category, and the color represents the mean of the strictly standardized mean difference (SSMD) of the significant assays in that category. Biological replicates from independent embryo batches are shown, comparing homozygous −/− to wildtype +/+ siblings except for duplicated a/b genes, which compare −/−;−/− to +/−;+/−. N for all experiments is available in Table S1. The comparisons of each genotype separately are available in Supplementary Fig. S13 and Supplementary Fig. S14. Baseline behavior: frequency of movement (e.g., number of bouts / hour), magnitude of movement (e.g., bout distance traveled), and location (e.g., fraction of bout in center of well) preferences were calculated for the baseline data. Stimulus-driven behavior: response frequency, latency, and magnitude for dark flashes (DF), light flashes, acoustic habituation (Hab), and acoustic stimulation that is strong, weak, or strong preceded by weak (PPI). **f**) Examples of *atp1a3a-ko* behavioral phenotypes. *p* values: non-significant for full time course (shown) but significant for most subsections, 6.2e-9 for bout time, 2.0e-8 for seizure-like movements, and 8.6e-6 for response to light flash response frequency during day of 5 dpf. **g**) Examples of *atp1a3a-pt* behavioral phenotypes. *p* values: 0.015 for bout frequency, 4.1e-5 for bout time. **h**) Examples of *atp1a3b-ko* behavioral phenotypes. *p* values: 0.0037 for bout frequency, 0.0036 for strong tap response frequency when preceded by a weak prepulse sound (sound frequencies 1400 Hz) during day of 5 dpf. Replicate data for f-h are available in Supplementary Fig. S10. Plots in f-h represent mean ± s.e.m. *p* values are Kruskal-Wallis ANOVA. The N for all experiments is available in Supplementary Table S1.

In contrast to the brain imaging findings, *atp1a3b* knockouts had stronger behavioral phenotypes than both *atp1a3a* mutants (**Fig. 2e**, **Supplementary Fig. S13**). *Atp1a3b-ko* had substantially reduced bout count, while *atp1a3a* movement frequency was elevated at night, in line with a previous zebrafish study linking the gene to sleep^40^. The *atp1a3a-pt* line exhibited similar, albeit milder, phenotypes than the *atp1a3a-ko* (**Fig. 2e**, **Fig. 2f**, **Fig. 2g**), whereas the point mutation in *atp1a3b* had almost no effect. The *atp1a3b-ko, atp1a3a-ko,* and *atp1a3a-pt* lines all displayed a preference for the center of the well, dramatic increases in the length of these movements (e.g., bout time) and a reduced responsiveness to dark flashes (**Fig. 2e**, **Supplementary Fig. S14**). Dark flash responses were most severely impacted in *atp1a3b-ko*, and both *atp1a3a* mutants displayed an increased frequency of responses to light flashes (**Fig. 2e**, **Fig. 2f**). Acoustic responses were relatively normal for both *atp1a3a* lines but substantially differed in *atp1a3b* knockouts (**Fig. 2h**). The response to weak acoustic stimulation was far lower, indicating an increased threshold to response, and the response to louder stimulation was enhanced, particularly in the prepulse inhibition paradigm when that loud sound was preceded by a weak prepulse. These different phenotypes are consistent with the expression patterns, given that *atp1a3b* is strongly localized to hindbrain circuits^40^, which control the acoustic startle response^41^.

From these SCHEMA and COS datasets, we selected the *atp1a3a-pt* and *sp4* mutants for additional investigation. Although its brain activity phenotype was mild, the *SP4* gene rose to the top of our list because of its location and because it is implicated in schizophrenia by GWAS^5^ in addition being a SCHEMA hit, further reinforcing its importance in this disorder.

### *Both atp1a3a* mutants possess juvenile spatial navigation deficit in Y-maze

Impairment in working memory is a core cognitive symptom of schizophrenia^42^. Because *sp4* and *atp1a3a* mutants both showed changes in activity of the forebrain, which is highly linked to working memory in both zebrafish and humans, we next probed working memory using the free movement pattern (FMP) Y-maze^43^ in juvenile animals. In this assay, zebrafish navigate a Y-maze, and their tetragrams, or four successive turns into arms of the Y-maze (left/L or right/R), are calculated. Zebrafish and other vertebrates exhibit a bias towards alternation tetragrams (LRLR or RLRL), and a higher percentage of alternation tetragrams compared to all tetragrams is a measure of working memory performance. We compared the percentage of alternation tetragrams and total number of tetragrams (as a proxy for activity) of the *sp4* mutant and both *atp1a3a* mutants to sibling controls^44,45^. The *sp4* mutant had no difference in alternation tetragram percentage compared to wildtype siblings (**Fig. 3a**). There was also a slight, but significant, increase in total tetragrams in *sp4* mutants. Both *atp1a3a* lines had lower alternation tetragram percentages compared to wildtype siblings (**Fig. 3b**, **Fig. 3c**). There were opposing effects on the total tetragram number, with the knockouts performing significantly fewer tetragrams than sibling controls, and the point mutants performing more tetragrams in more of the time bins than controls. This effect was driven by the second replicate (**Supplementary Fig. S15**). The reduced tetragrams observed in the knockouts, which suggest reduced activity, are consistent with poor health of the fish. In spite of this, the reduced alternation percent tetragrams in both mutants suggest that *atp1a3a* is required for working memory.

**Fig. 3:**
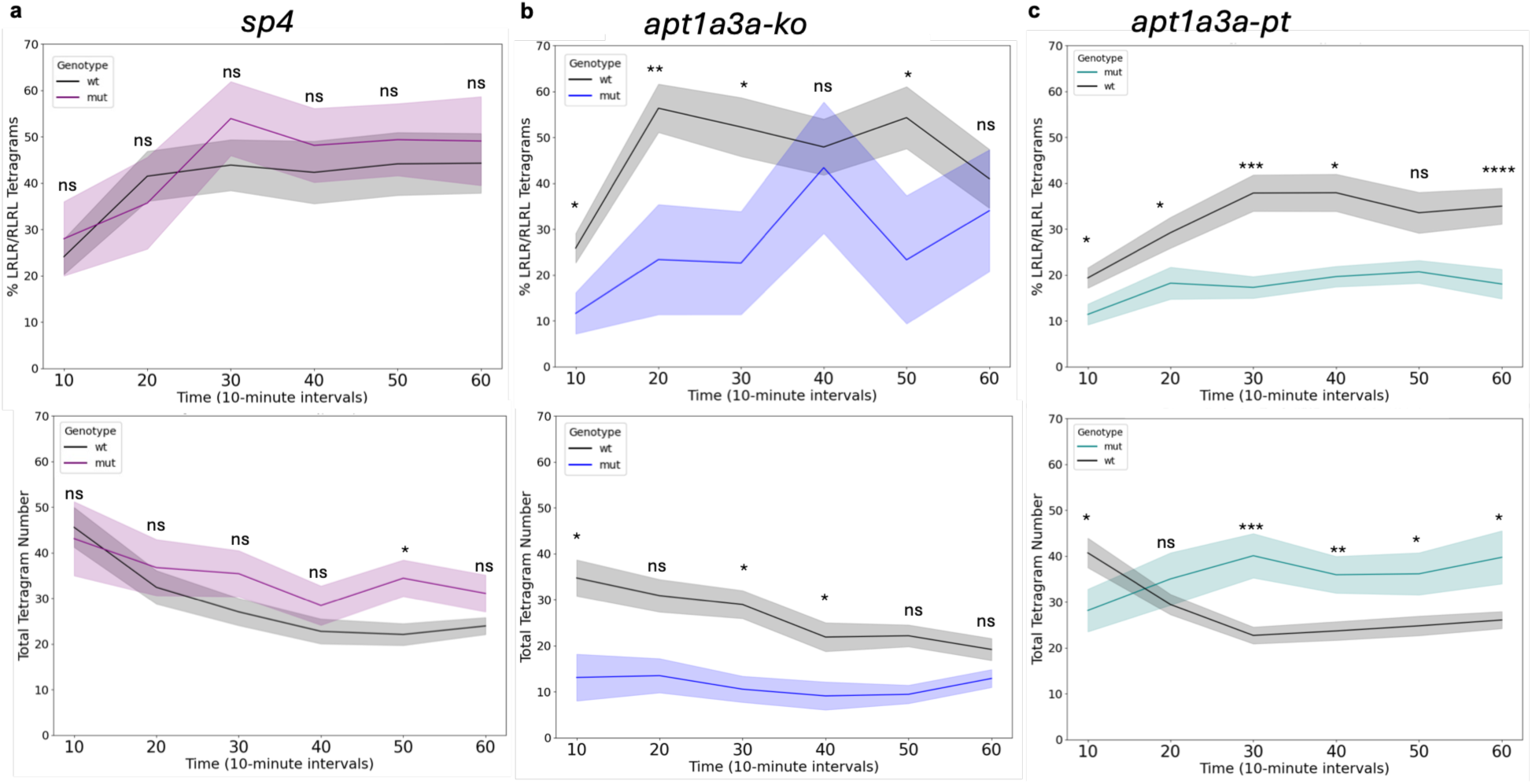
Juvenile behavioral phenotypes in a Y-maze assay. Percentage of alternation (%LRLR/RLRL) tetragrams (top) and total number of tetragrams performed (bottom) by *sp4* (**a**), *apt1a3a-ko* (**b**), and *atp1a3a-pt* (**c**) zebrafish. Means and standard error are plotted at each time point, and each point represents a bin of the previous ten minutes. Significance denotes pairwise comparisons between wildtype (black) and mutant (other) fish at that individual time point, performed after a Two-Way ANOVA (ns= not significant, * *p*<0.05, ** *p*<0.01, ** *p*<0.001, **** *p*<0.0001). Genotype main effect results of Two-Way ANOVA for each gene: Alternation Percent – *sp4 p*= 0.36; *atp1a3a-ko p*=0.00011; *atp1a3a-pt p*<0.0001; Total Tetragrams – *sp4 p*=0.016, *atp1a3a-ko p*<0.0001, *atp1a3a-pt p*=0.00014).

### Sterol biosynthesis genes are upregulated in adult *atp1a3a-pt* mutant brains

To probe the molecular changes underlying brain activity and Y-maze phenotypes, we performed bulk RNA-seq of 6 dpf heads from *atp1a3a-pt* mutants and wildtype siblings (**Supplementary Fig. S16a-b**, **Supplementary Table S2**). Gene set enrichment analyses (GSEA) revealed that differentially expressed genes were dominated by markers of the endocrine pancreas, where *atp1a3a* is also expressed^46^ (**Supplementary Fig. S16c-d**).

To avoid contamination from non-neural tissue, we next conducted bulk RNA-seq on dissected adult brains. Differential gene expression analysis revealed relatively few dysregulated genes, with 53 upregulated genes and 71 downregulated genes at a threshold of padj < 0.05 (**Fig. 4a**, **Supplementary Table S3**). Genes encoded by chromosome 19, where the *atp1a3a* gene is located, were significantly overrepresented among dysregulated genes (**Supplementary Fig. S17a**). GSEA using gene sets derived from previously published forebrain single-cell data showed negative enrichment of the Progenitor_02 cluster (**Supplementary Fig. S17b**) and positive enrichment of the Oligodendrocyte cluster (**Supplementary Fig. S17c**), although the Oligodendrocyte cluster enrichment was largely driven by a single mutant sample.

**Fig. 4:**
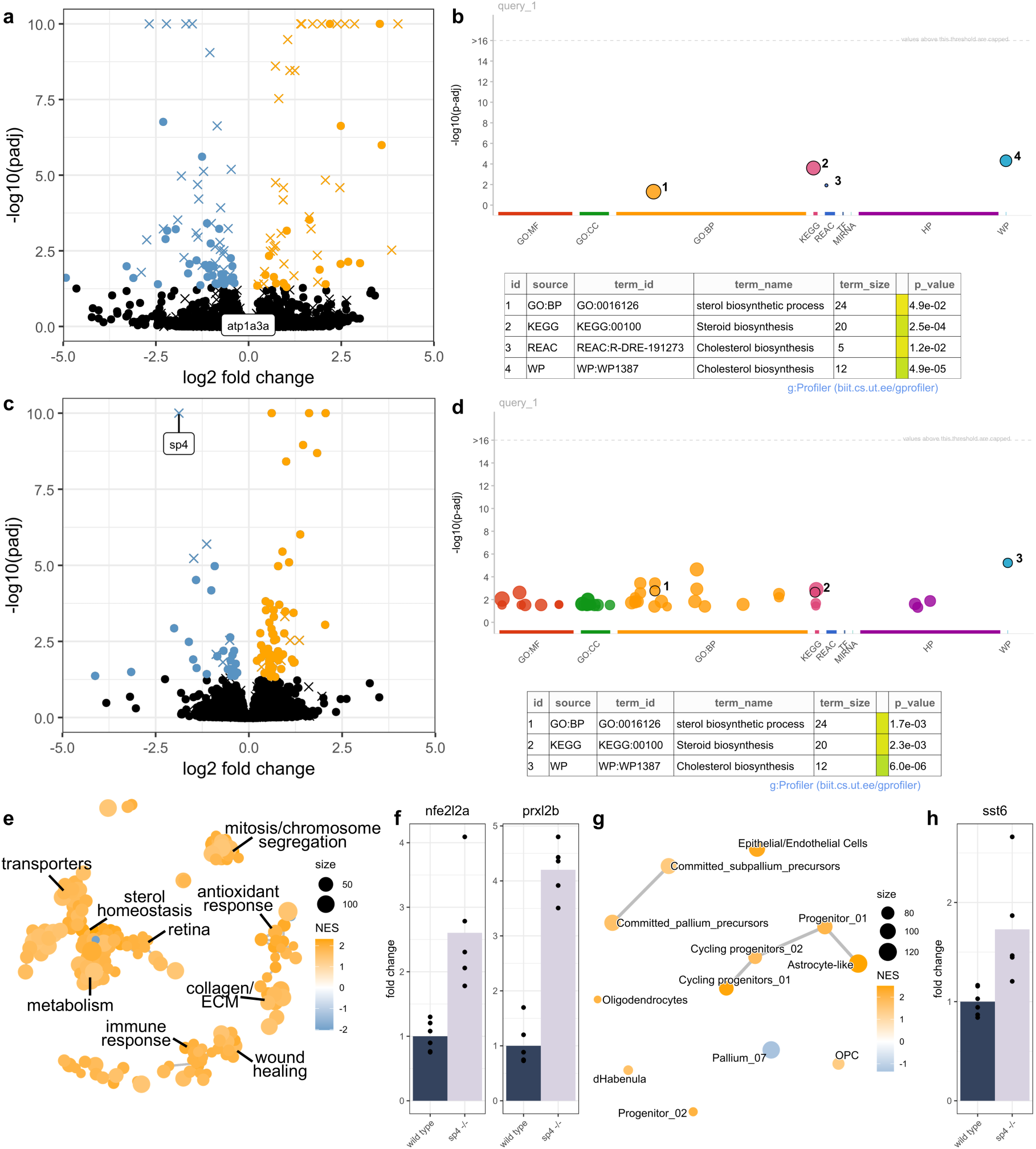
Bulk RNA-seq in *atp1a3a-pt* and *sp4* adult brains highlights shared and divergent transcriptomic phenotypes. **a**) Volcano plot displaying differentially expressed genes in homozygous *atp1a3a-pt* adult brains compared to wildtype siblings. Blue indicates downregulated genes and orange indicates upregulated genes at a threshold of padj < 0.05. Genes encoded by zebrafish chromosome 19 are denoted with the X symbol. −log10(*padj*) values are capped at 10 for display. **b**) Manhattan plot representing significant terms identified by GO analysis of upregulated genes in *atp1a3a* mutants. Point size denotes size of the term. **c**) Volcano plot displaying differentially expressed genes in homozygous *sp4−/−* adult brains compared to wildtype controls. Blue indicates downregulated genes and orange indicates upregulated genes at a threshold of *padj* < 0.05. Genes encoded by zebrafish chromosome 19 are denoted with the X symbol. −log10(*padj*) values are capped at 10 for display. **d**) Manhattan plot representing significant terms identified by GO analysis of upregulated genes in *sp4* mutants. Only select terms related to sterol biosynthesis are highlighted. Point size denotes size of the term. **e**) Network plot of significantly enriched terms in *sp4−/−* mutants from GSEA with C5 gene sets from the Molecular Signatures Database. Nodes represent enriched terms and edges join terms with overlapping genes. **f**) Upregulation of *nfe2l2a* (*p* value, *padj* = 4.08E-10, 9.67E-07) and *prxl2b* (*p* value, *padj* = 3.23E-21, 2.81E-17) in *sp4−/−* mutants compared to control. **g**) Network plot of significantly enriched terms from GSEA with gene sets derived from previously published forebrain single-cell data. Nodes represent enriched terms and edges join terms with overlapping genes. **h**) Upregulation of *sst6* (*p* value, *padj* = 4.46E-05, 0.013) in *sp4−/−* mutants compared to control.

Gene ontology (GO) analysis of upregulated genes identified terms related to sterol biosynthesis, with overexpression of genes including *cyb5r2*, *cyp51*, *hsd17b7*, *lss*, and *nsdhl* (**Fig. 4b**). At a relaxed threshold of unadjusted *p* < 0.05, several genes with potential links to schizophrenia risk showed the same direction of effect in 6 dpf and adult samples (*dlg1*, *il21r.1*, *map2k7*) (**Supplementary Fig. S18a**). In contrast, the immediate early gene *fosaa* was downregulated only at 6 dpf (**Supplementary Fig. S18b**), and *prkd1*, *rims1b*, and *rims2a*, two of which are orthologs of schizophrenia risk genes^5^, were downregulated only in adult samples (**Supplementary Fig. S18c**). Together, these analyses reveal relatively modest changes in the transcriptome of *atp1a3a-pt* samples, with differential expression of individual genes involved in synaptic structure and transmission, sterol synthesis, and immune function.

### Loss of *sp4* dysregulates glial and metabolic transcripts in the adult brain

We next performed bulk RNA-seq on mutant *sp4−/−* adult brains and wildtype controls. At a threshold of padj < 0.05, we identified 66 upregulated genes and 33 downregulated genes compared to control (**Fig. 4c**, **Supplementary Table S4**). Like *atp1a3a*, *sp4* is encoded by zebrafish chromosome 19, and we observed a significant enrichment of differentially expressed genes on chromosome 19 (**Supplementary Fig. S19a**). GO analysis of upregulated genes identified enrichment of Notch signaling as well as several terms related to sterol biosynthesis and metabolism (**Fig. 4d**).

GSEA using C5 gene sets from the Molecular Signatures Database highlighted positive enrichment of terms related to mitosis, antioxidant response, and metabolic processes including sterol and fatty acid synthesis (**Fig. 4e**, **Supplementary Fig. S19b-c**). Top genes contributing to oxidant detoxification and fatty acid metabolism were *nfe2l2a* (*nrf2*) and *prxl2b*, respectively (**Fig. 4f**). GSEA using terms from forebrain single-cell data showed positive enrichment of multiple radial glial, astrocyte-like, and oligodendrocyte subtypes and negative enrichment of Pallium_07 (**Fig. 4g**, **Supplementary Fig. S19d**). Overexpression of the neuropeptide *sst6* (**Fig. 4h**) contributed to enrichment of dHabenula, which is a brain region that has been linked to schizophrenia in humans and in mouse models^47^.

### Shared cholesterol synthesis phenotype in *atp1a3a-pt* and *sp4* mutant transcriptomes

We observed that cholesterol synthesis genes such as *msmo1* and *sqlea* contributed to enrichment of both astrocyte-like and sterol synthesis terms in GSEA of *sp4* mutant data, which suggests that cholesterol synthesis genes may serve as marker genes for astrocyte-like cells (**Supplementary Fig. S19c-d**). In the adult mammalian brain, astrocytes produce and release cholesterol for use by neurons^48^. To explore the cell-type specific expression of cholesterol synthesis genes across the zebrafish lifetime, we re-analyzed previously published single-cell RNA-seq data collected from the zebrafish telencephalon at 6 dpf, 15 dpf, and adult stages^49^. We first merged clusters to generate a *ptn+/her4+/dla-* astrocyte-like cluster, a *ptn-/her4+/dla+* radial glial cluster, and a *snap25a+* neuronal cluster (**Supplementary Fig. S20a-b**). The astrocyte-like cluster expresses the markers *s100b and gfap* and represents a heterogeneous population of cells that has variably been called radial astroglia, quiescent radial glia, and astrocytes^50^. Expression of cholesterol synthesis genes, including *msmo1* (**Supplementary Fig. S20c**), showed increased expression in astrocyte-like cells compared to radial glia and neurons, and expression in astrocyte-like cells increased from 6 dpf to 15 dpf (**Supplementary Fig. S20d**). This analysis suggests that telencephalic astrocyte-like cells express the genes required to synthesize cholesterol and that cell-type-specific expression of these genes changes across the zebrafish lifetime.

Mammalian astrocytes are thought to use the Bloch pathway for cholesterol synthesis^51^ (**Fig. 5a**), and examination of cholesterol synthesis, metabolism, and regulatory genes in *atp1a3a-pt* (**Fig. 5b**) and *sp4* (**Fig. 5c**) adult RNA-seq data showed upregulation of some but not all members of the cholesterol synthesis pathway. For example, *hsd17b7*, *msmo1*, and *srebf2* were upregulated in both *atp1a3a-pt* and *sp4* mutant brains compared to control (**Fig. 5d**). SREBF2 is the major transcription factor regulating the cholesterol synthesis pathway^52^, and its loss in astrocytes alters mouse brain development and behavior^53^. Several genes in schizophrenia risk loci have roles in cholesterol and lipid metabolism (**Supplementary Fig. S21**), including *MSMO1* (**Fig. 5e**) and *SREBF2*, suggesting that changes to expression of genes related to cholesterol metabolism can directly contribute to disorder susceptibility.

**Fig. 5:**
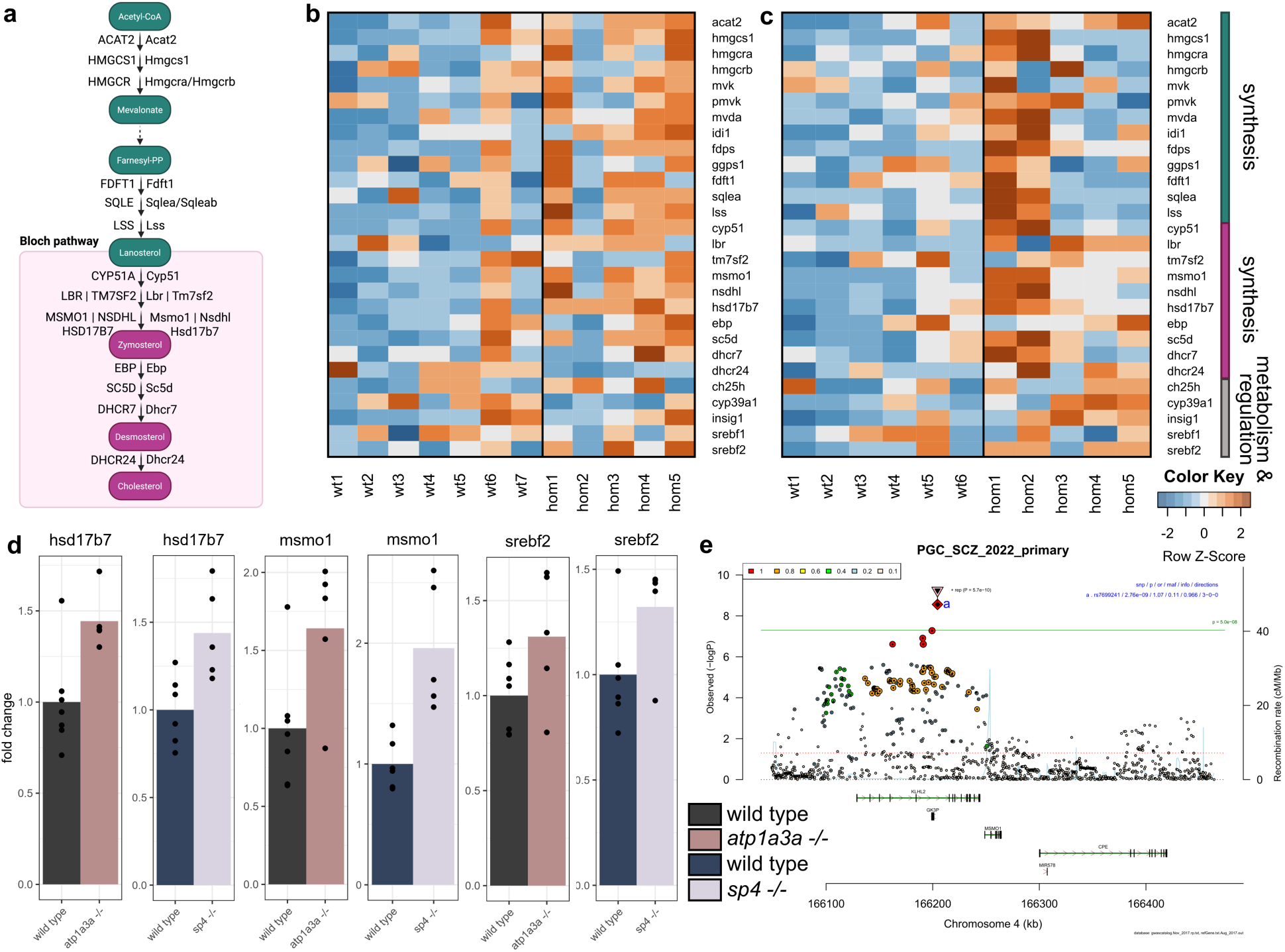
Dysregulation of cholesterol synthesis pathway genes in *atp1a3a-pt* and *sp4* mutant brains. **a**) Simplified cholesterol synthesis pathway, modified from Wikipathways (WP4346). The Bloch pathway is thought to be the predominant synthesis pathway in mammalian astrocytes. Human protein names are on the left and putative zebrafish orthologs are on the right. **b**) Heatmap showing relative expression of cholesterol synthesis genes in *atp1a3a-pt −/−* brains compared to controls. **c**) Heatmap showing relative expression of cholesterol synthesis in *sp4 −/−* brains compared to controls. **d**) Upregulation of *hsd17b7*, *msmo1*, and *srebf2* in *atp1a3a* and *sp4* brains. Raw and adjusted *p* values are in Supplementary Tables S3 and S4. **e**) RICOPILI plot for associated locus containing *MSMO1* from the 2022 PGC study.

### Abnormal free cholesterol and Fabp7a staining in juvenile *atp1a3a* and *sp4* mutant brains

To validate findings from our bulk and single-cell RNA-seq analyses, we stained 21 dpf brains for markers of cholesterol, cholesterol synthesis, and glia. In wildtype animals, hybridization chain reaction (HCR) labeling of the *msmo1* transcript co-localized with the astrocyte-like/radial glial marker Fabp7a **(Fig. 6a)** along the midline of the telencephalon, confirming our single-cell RNA-seq finding that glia express this component of the cholesterol biosynthesis pathway. We next stained *atp1a3a-ko*, *atp1a3a-pt*, and *sp4* mutant brains using filipin, which labels free cholesterol **(Fig. 6b)**. All three mutants showed a significant increase in filipin staining throughout the brain compared to wildtype controls **(Fig. 6c)**. Both *atp1a3a-pt* and *atp1a3a-ko* brains were significantly smaller than wildtype siblings, with *atp1a3a-ko* brains having less than half of the area of wildtype siblings in a z-projection **(Fig. 6d)**. We next sought to determine whether the changes in free cholesterol in *atp1a3a-pt* and *sp4* mutants were localized to astrocyte-like cells by staining juvenile brains using a Fabp7a antibody **(Fig. 6e)**. However, despite no difference in *fabp7a* transcript levels in adult RNA-seq data, both mutants showed a significant reduction of Fabp7a staining in the midline of the forebrain at 21 dpf **(Fig. 6f)**, suggesting potential age-dependent changes in *fabp7a* expression, post-translational regulation of the Fabp7a protein, or alterations in localization/antibody accessibility. We also observed disorganization of telencephalic structures in the *sp4* mutants compared to control **(Fig. 6g)**.

**Fig. 6:**
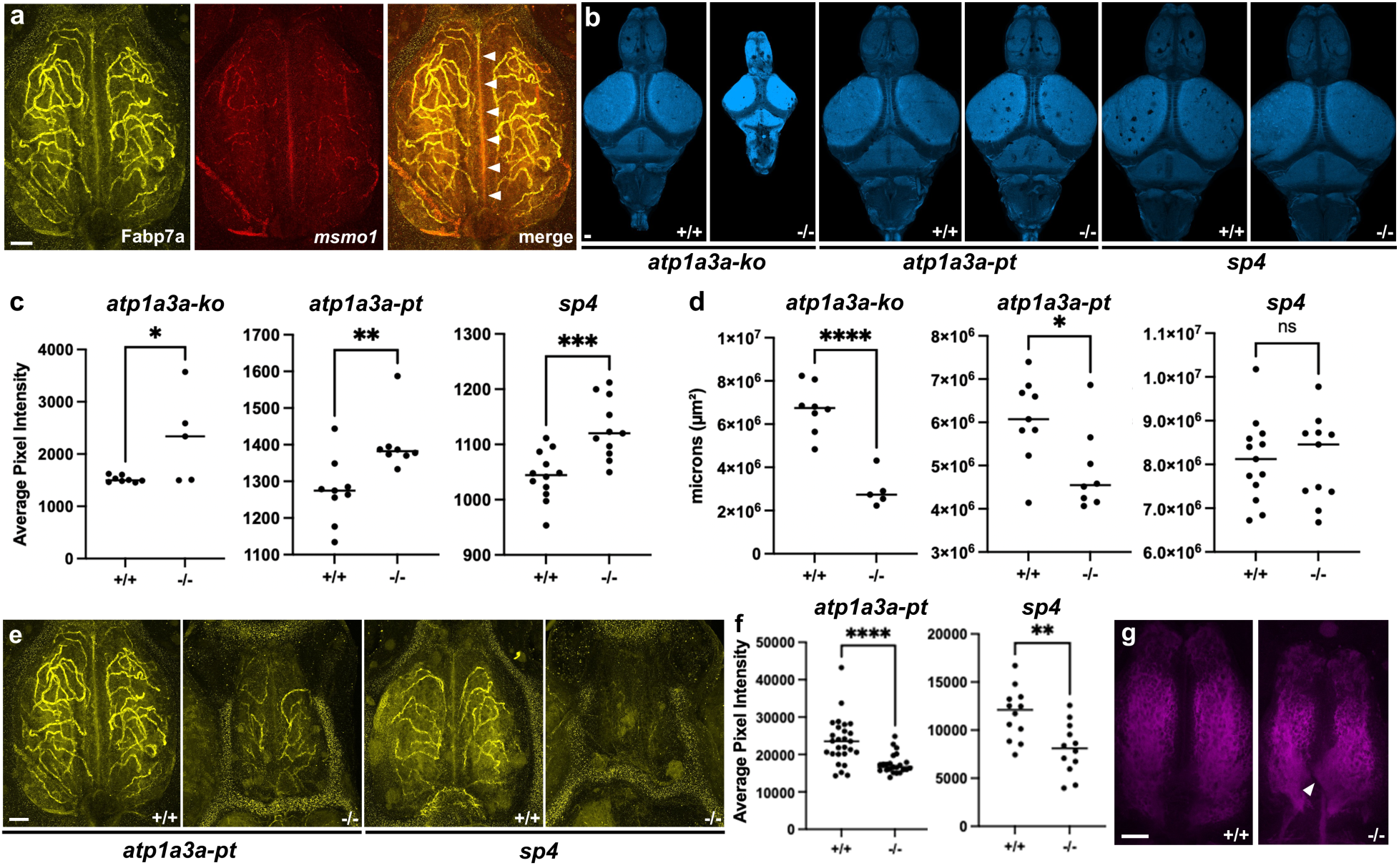
Aberrant free cholesterol and glial staining in *atp1a3a* and *sp4* juvenile brains. **a)** Fabp7a immunostaining (left), *msmo1* HCR (center), and merged maximum intensity z-projection (MIP) of wildtype telencephalon at 21 dpf. Arrowheads denote colocalization of signal along the midline. **b)** Representative filipin staining of *atp1a3a-ko*, *atp1a3a-pt*, and *sp4* homozygous mutant and wildtype sibling brains at 21 dpf. (MIP) **c)** Quantification of filipin staining based on mean intensity values from sum-of-slices projections. Unpaired t-tests showing increased filipin staining for *atp1a3a-ko* (*p* = 0.0238), *atp1a3a-pt* (*p* = 0.0055), and *sp4* (*p =* 0.0005). **d)** Quantification of brain total area (µm²) based on sum-of-slices projections of filipin staining. Unpaired t-tests where *atp1a3a-ko* (*p* < 0.0001), *atp1a3a-pt* (*p* = 0.0238), and *sp4* (ns). **e)** Representative Fabp7a immunostaining in the telencephalon of 21 dpf *atp1a3a-pt* and *sp4* mutants and wildtype controls. (MIP) **f)** Quantification of Fabp7a immunostaining along the midline of the (MIP). Unpaired t-tests show reduced Fabp7a staining for *atp1a3a-pt* (*p* < 0.0001) and *sp4* (*p =* 0.0032). **g)** MIP z-projection of representative tERK staining in 21 dpf *sp4* mutant and wildtype sibling telencephalon. Arrowhead denotes example of dysmorphology. For all panels, scale bar = 50 µm and ns= not significant, * *p*<0.05, ** *p*<0.01, ** *p*<0.001, **** *p*<0.0001.

## Discussion

Here, we showed that mutations in two high-confidence schizophrenia risk genes converge on dysregulation of sterol metabolism in the adult zebrafish brain. We first generated zebrafish lines representing several types of schizophrenia genetic risk (**Supplementary Table S1**) and identified altered brain activity and behavior in knockouts of genes implicated by large-scale exome sequencing efforts, as well as a rare *de novo* variant likely causative of childhood-onset schizophrenia. RNA-sequencing of two prioritized lines, a missense mutation in *atp1a3a* and protein-truncating allele of *sp4*, revealed convergent dysregulation of cholesterol metabolism in the brain. Free cholesterol was elevated in both lines, and Fabp7a, which is involved in transport of fatty acids and marks astrocyte-like cells in the zebrafish telencephalon, was downregulated (**Fig. 6**). The convergence of shared phenotypes between *sp4* and *atp1a3a* mutants raises a central question: does sterol dysregulation represent a primary pathogenic mechanism or a secondary response to altered neuronal activity?

One possibility is that sterol pathway activation arises as a downstream consequence of altered neuronal excitability and metabolic stress. Synaptic cholesterol is required for plasticity and synapse growth and responds to neuronal activity^54–56^. *Sp4* itself is regulated by neuronal activity: its knockdown in cerebellar granule neurons blocks dendritic remodeling in response to depolarization^57^, possibly through a phosphorylation event downstream of NMDA receptor activation^58^. Neuronal activity also generates oxidative stress, activating astrocytic NRF2 (NFE2L2)^59^, a master regulator of antioxidant response. Intriguingly, *nfe2l2a* is one of the most upregulated genes in the *sp4* zebrafish knockouts, and it directs metabolic reprogramming and lipid homeostasis in response to stress^60–62^. Together, alterations in the sterol pathway in an activity-dependent transcription factor (*sp4*), a Na+/K+ ATPase pump (*atp1a3a*), and a previously published subunit of the NMDA receptor (*Grin2a*) suggest that aberrant neuronal activity may evoke changes in cholesterol demand or synthesis. Defining the causal relationships between activity, stress, and metabolism will be essential for determining appropriate clinical interventions. Epidemiological studies of statins, which can reduce cholesterol in the brain depending on the compound, have been associated with improvement in psychiatric symptoms, hospitalization, and suicide risk^63–67^. To evaluate whether sterol dysregulation represents a primary genetic vulnerability rather than a purely compensatory response, we next consider evidence from human genetic studies.

Both common variants identified via GWAS and rare coding variants implicate alterations to cholesterol and lipid metabolism in schizophrenia risk. In addition to *MSMO1* and *SREBF2*, schizophrenia GWAS risk loci contain numerous genes directly involved in these pathways (**Supplementary Fig. S21**, **Table 1**). Most of the genes listed in Table 1 are located in loci with relatively few genes, increasing the likelihood that they drive the association signal. In addition to being a GWAS hit, *FAM120A* was in the top 20 SCHEMA genes^23^. Another SCHEMA gene, *CUL1*, is part of the ubiquitin ligase complex directly responsible for the degradation of multiple SREBP transcription factors^68^. Beyond the top SCHEMA genes, *FABP7* and *HMGCR* also carry protein-truncating variants with odds ratios of 6.02 (*p* value 0.000553) and 4.46 (*p* value 0.00197), respectively. Among recurrent CNVs of large effect, *MLXIPL*^69^ is one of ~25 genes in the 7q11.23 duplication region, and the minimal 16p11.2 distal deletion^70^ includes *SPNS1*, whose deficiency leads to lysosomal cholesterol accumulation and impaired lipid metabolism^71,72^. In zebrafish adult brains, *spns1* is enriched in astrocyte-like cells^49^. Further reinforcing this connection, at least 25% of individuals with Niemann-Pick disease type C, caused by improper cholesterol trafficking leading to its accumulation^73^, develop schizophrenia-like psychosis preceding neurodegeneration^74–77^. Collectively, these heterogeneous findings point to glial control of neuronal lipid and cholesterol supply as a potential recurrent vulnerability node across diverse forms of genetic risk.

**Table 1:**
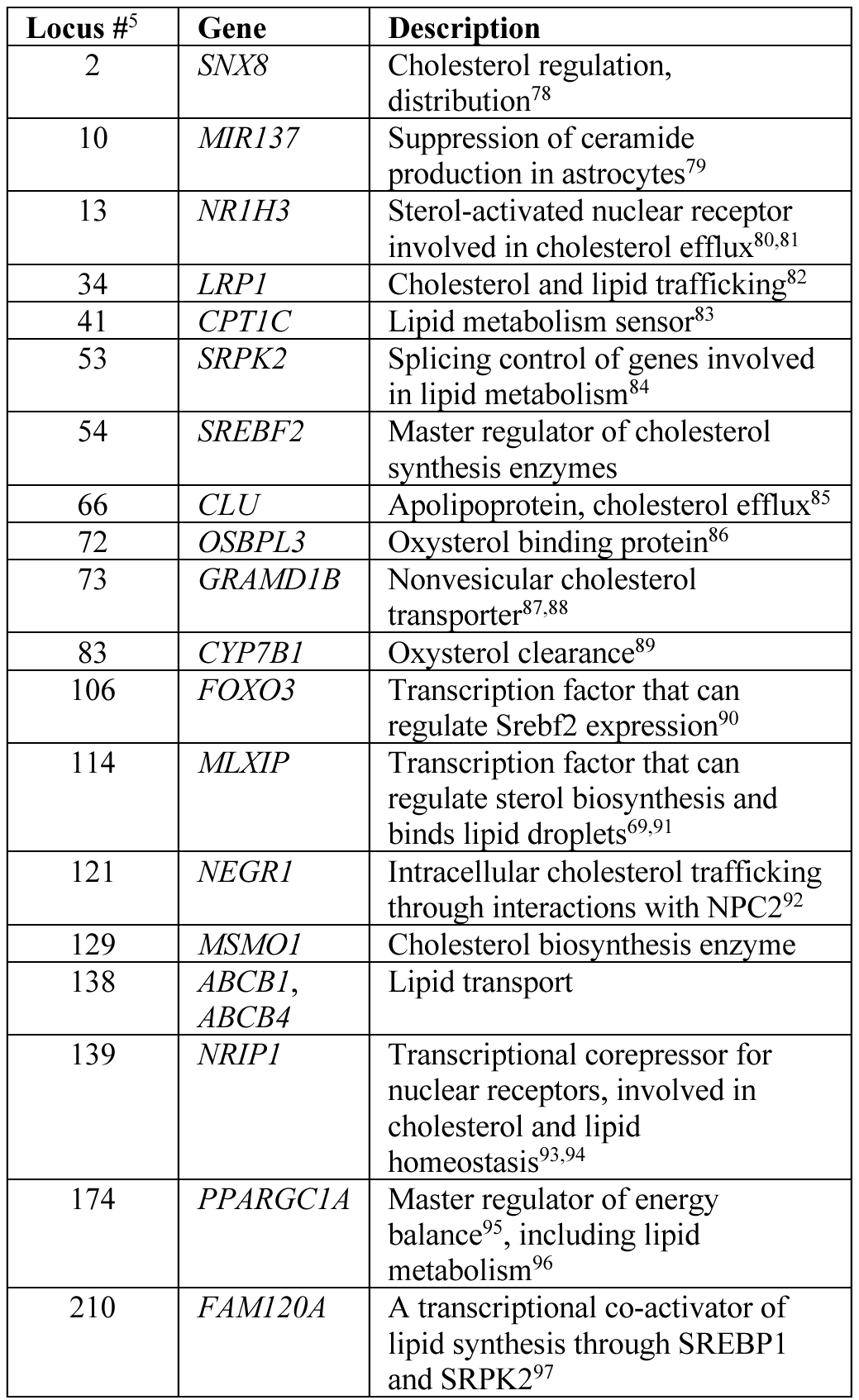
Selected genes involved in cholesterol and lipid metabolism and located in schizophrenia GWAS loci from the 2022 PGC study.

Astrocyte-derived cholesterol production is dynamically regulated across development and aging and is required for synapse formation and stabilization^56^. In the postnatal mammalian brain, neurons downregulate cholesterol synthesis and astrocytes become the primary source. Our re-analysis of zebrafish single-cell RNA sequencing data identified conservation of this age-related shift, as expression of core cholesterol synthesis genes increased in astrocyte-like cells between larval and juvenile stages (**Supplementary Fig. S20**). Interestingly, although core sterol synthesis genes were upregulated in both *atp1a3a-pt* and *sp4* mutants, Fabp7a immunoreactivity was reduced. Fabp7a is a lipid-binding protein associated with astrocyte-like cells, and its downregulation may reflect altered fatty acid handling or a shift in astrocyte state rather than a simple loss of glial identity. Sterol synthesis is also relevant in human tissue: single-nucleus RNA sequencing (snRNA-seq) revealed decreased expression of genes involved in cholesterol and fatty acid synthesis and export in the postmortem prefrontal cortex of individuals with schizophrenia^14^. This decrease contrasts with our findings of increased RNA levels and free cholesterol. Either direction of transcriptional change could reflect compensation, and snRNA-seq could undercapture transcripts localized to peripheral astrocytic processes. Recently, mouse knockouts for two SCHEMA genes, *Akap11* and *Grin2a*, both showed significant transcriptional upregulation of sterol and lipid synthesis pathways in astrocytes^98,99^. These changes were developmentally regulated, as they were not significant at 4 weeks in *Grin2a* mutants but emerged at 12 weeks. Cholesterol levels were not directly assessed in the *Grin2a* line, but *Akap11* deficiency led to accumulation of cholesteryl esters and lipids in isolated astrocytes and brain tissue. Further complicating interpretation, although transcription of cholesterol metabolism genes is upregulated in *Grin2a* mutants, a recent preprint suggests that their translation is downregulated^100^. These findings are not yet reconciled but collectively indicate dysregulation of sterol gene programs. Their developmental regulation is intriguing given the high demand for cholesterol during adolescent myelination and synaptic remodeling, which coincides with the typical timeline of schizophrenia pathogenesis. These cell-type-specific developmental dynamics may help explain why convergence across risk genes emerges preferentially at later stages.

Our findings also underscore the value of incorporating rare, high-effect variants when developing functional models of schizophrenia. Although COS is rare, such rare variants may provide insight into general forms of schizophrenia. For example, the protein encoded by *ATP1A3*, which is mutated in multiple individuals with COS, is downregulated in the auditory cortex of individuals with a more typical schizophrenia progression^101^. Large-scale population genetic studies suggest that rare coding variant analyses tend to prioritize more trait-specific genes than GWAS^102^, potentially enriching for schizophrenia-relevant biology. In prioritizing *atp1a3a-pt* and *sp4*, our study revealed that convergent phenotypes may only emerge at later developmental stages. Since convergence may be stage dependent, mutants lacking larval phenotypes in this study or our previous work^21^ should not be considered dead ends. Future zebrafish studies should examine multiple stages and include rare, high-effect variants for mechanistic entry points, even if they lack strong population-wide support. Zebrafish are particularly well suited for longitudinal analysis, providing an opportunity to determine how cholesterol synthesis is regulated in response to neuronal activity and in specific cell types across the lifespan.

## Materials and Methods

### Zebrafish husbandry

All zebrafish procedures were approved by the UAB Institutional Animal Care and Use Committee (IACUC protocols 22155 and 21744) and UMass Chan Institutional Animal Care and Use Committee (IACUC protocol 202300000053). Mutant lines were generated in an Ekkwill-derived background using CRISPR/Cas9 genome editing as previously reported^21,25^. Mutant sequences, guide RNA targets, and genotyping primers are available in Supplementary Table S1. Adult fish and larvae were housed at 28°C under a 14-hour light/10-hour dark cycle. Experimental larvae were raised in 150 mm Petri dishes containing fish water with methylene blue at densities below 160 larvae per dish. Debris was removed at least twice before brain imaging and behavior experiments. Larvae lacking inflated swim bladders were excluded. Control animals consisted of siblings from a single breeding pair, and all individuals were genotyped following completion of experiments.

### Brain activity and morphology

Immunostaining for phosphorylated ERK (pErk) was performed according to previously published methods^21,25,37^. All samples were collected in the afternoon of 6 dpf. The total ERK antibody (Cell Signaling, #4696) was used at a 3:1000 dilution, and the pErk antibody (Cell Signaling, #4370) was used at 1:500. Primary antibody incubation typically lasted 2–3 days, followed by a one-day incubation with secondary antibodies. Confocal image stacks were acquired using a Zeiss LSM 900 upright microscope equipped with a 20×/1.0 NA water-dipping objective. Image volumes were aligned to the standard zebrafish Z-Brain reference atlas using the Computational Morphometry Toolkit (CMTK)^37,103,104^. Statistical analysis with MapMAPPING^21,37^ employed a false discovery rate (FDR) threshold calibrated such that 0.05% of control pixels were designated significant for both neural activity and structural comparisons. Sample sizes for each assay are listed in Supplementary Table S1.

### Juvenile *in situ* hybridization chain reaction (HCR) and antibody staining

At 21 dpf, zebrafish were fixed overnight at 4°C in 4% paraformaldehyde (PFA), followed by three 5-minute washes in PBST. After fixation larvae were genotyped, and homozygous mutant and wildtype siblings were selected for HCR. Homozygous mutants were tail clipped to mix with wildtype siblings. A *msmo1* probe set (available on Zenodo) was generated using a publicly available probe design pipeline (https://github.com/rwnull/insitu_probe_generator). HCR was carried out on 21 dpf zebrafish following the Molecular Instruments RNA FISH v3.0 protocol, with minor modifications as described. Following rehydration, larvae were depigmented in a solution containing 3% hydrogen peroxide and 0.8% potassium hydroxide for 7-8 minutes. Samples were then incubated in Proteinase K (Takara, #9034) for 30 minutes at room temperature to enhance probe penetration, followed by three washes (5 minutes each) in PBST at room temperature. For detection of *msmo1* transcripts, 4 pmol of the designed probe set was applied to each sample. Signal amplification was performed using B1-548 H1 and H2 hairpins (3 μM stock), which were snap-cooled according to the manufacturer’s instructions. A total of 2 μL of each hairpin was diluted into 150 μL of amplification buffer and added to the samples for overnight incubation at room temperature. For simultaneous detection of Fabp7a, the Fabp7 antibody (Millipore Sigma, ABN14, 1/500) was included during the amplification step. After amplification, samples were washed twice for 5 minutes, once for 1 hour and once for 5 minutes in 5x SSCT at room temperature. The larvae were then mounted in low-melt agarose and imaged using confocal microscopy.

### Filipin staining of juvenile zebrafish brains

At 21 dpf, zebrafish were fixed overnight at 4°C in 4% PFA, followed by three 5-minute washes in 1× PBST. After fixation, larvae were genotyped, and homozygous mutant and wildtype siblings were selected for brain dissection. Filipin staining was performed as previously described^105^ with minor modifications. Dissected brains were incubated in Filipin III solutions (Sigma-Aldrich) for 1 hour at room temperature, followed by overnight incubation at 4°C and an additional 3-hour incubation at room temperature, rotating and protected from light. Following staining, brains were washed three times for 5 minutes each in 1× PBST. Samples were then mounted and imaged using confocal microscopy.

### Analysis of juvenile imaging data

Image analysis was performed using Fiji (ImageJ)^106^. For each z-stack, both maximum intensity projection (MIP) and sum of slices projection images were generated. Regions of interest (ROIs) were defined on the MIP images to minimize background signal and ensure consistent anatomical boundaries across samples. For Fabp7a analysis, the ROI was restricted to the midline region of the forebrain, corresponding to the localization of astrocyte-like cells and the telencephalon domain of *msmo1* expression. After ROI selection, image channels were separated, and two-dimensional measurements were performed on the cropped MIP images to quantify mean fluorescence intensity of the Fabp7a signal. For filipin staining, two-dimensional measurements were performed on the cropped whole brain MIP images to generate binary masks corresponding to the selected ROIs. These masks were subsequently applied to the corresponding sum-of-slices projections, and mean fluorescence intensity and total area (µm²) were measured from the masked regions. Mean intensity values (sum-of-slices for filipin; MIP for Fabp7a) and total area measurements for filipin were imported into GraphPad Prism for statistical analysis. Normality was assessed using the Shapiro-Wilk test. Comparisons between homozygous mutant brains and wildtype sibling controls were performed using an unpaired t-test, with statistical significance defined as *p* < 0.05.

### Larval behavior

Behavioral experiments were carried out and analyzed as described previously^25^, using custom-built behavioral systems^107^. The assay begins on the evening of 4 days post-fertilization (dpf) and proceeds through 6 dpf. On 5 dpf, larvae were exposed to light flashes (9:11–9:25), mixed acoustic stimuli consisting of prepulse, strong, and weak sounds (9:38–2:59), and three blocks of acoustic habituation (3:35–6:35). During the intervening night (between 5 and 6 dpf), additional mixed acoustic stimuli (1:02–5:00) and light flashes (6:01–6:20) were delivered. On 6 dpf, larvae underwent three dark-flash blocks separated by one-hour intervals (10:00–3:00), followed by mixed acoustic stimuli and light flashes (4:02–6:00). Stimulus-driven responses were quantified from 1-second recordings captured at 285 frames-per-second (fps). Baseline locomotor metrics were quantified from positional tracking at approximately 30 fps and included movement frequency (e.g., active seconds per hour), bout magnitude (e.g., bout velocity), and spatial preference (e.g., proportion of bout time spent in the center of the well). For stimulus-evoked responses, frequency, response magnitude, and response latency were calculated from the high-speed recordings. Stimulus epochs were also subdivided for analysis (for example, early versus late dark flashes within a block). Behavioral analysis code is available at https://github.com/thymelab/ZebrafishBehavior. Sample sizes for each assay are listed in Supplementary Table S1.

A wide range of behavioral features was computed, and both raw and summarized measurements are presented. Summary visualizations were generated by grouping related measures (e.g., all frequency-based metrics). In these bubble plots, bubble size represents the percentage of significant measurements within a category, and color indicates the mean strictly standardized mean difference (SSMD) of significant assays in that group. In some cases, two offset bubbles may appear within a single category to reflect bidirectional effects within a merged measure (e.g., increases shown in purple and decreases in orange). This can occur when phenotypes vary across experimental timepoints, such as differences in locomotor activity between the night of 4 dpf and 6 dpf. Analysis scripts for data merging and bubble plot generation are available at https://github.com/thymelab/DownstreamAnalysis.

### Juvenile Y-maze assay

Y-maze assays were performed as previously described^45^. Briefly, we 3D-printed plates of 12 Y-mazes using ABS to test multiple fish in parallel. Single zebrafish were placed in a single arm of individual Y-mazes, with the rest of the maze blocked off using an ABS printed divider. After a 10-minute habituation period, the divider was lifted, and the fish was allowed to explore the Y-maze for one hour freely. Movies were captured using the FLIR FlyCapture SDK software. The same animal behavior box used in larval assays was used to record Y-maze behavior. We then used our custom tracking software, *StrIPETrack*^45^, to quantify turn direction, which was used to calculate total tetragrams and the percentage of alternation tetragrams.

### RNA-seq sample collection

Bulk RNA sequencing libraries were generated using a modified SMART-Seq2 workflow^108^ and as previously described^25^. Larval zebrafish heads were dissected following anesthesia, flash-frozen, and stored at −80°C, with corresponding bodies retained for genotyping. For each condition, four heads were pooled per biological replicate (3-5 replicates total). Additionally, adult brains were dissected following anesthesia, stored and genotyped similar to the larval heads. Adult brains were processed separately for downstream assays. RNA was isolated using E.Z.N.A. MicroElute Total RNA Kit (Omega Bio-Tek R6834-02) with DNase treatment, and concentration and purity were assessed spectrophotometrically.

For first-strand cDNA synthesis, 6 µL of extracted RNA was combined with 0.3 µL of 10 µM reverse transcription oligo (5′-AGACGTGTGCTCTTCCGATCT(30)VN-3′), 3 µL of 10 mM dNTP mix (Thermo Fisher, R0192), 0.3 µL RNase inhibitor (40 U/µL; Life Technologies, AM2694), and 2.5 µL of 1 M trehalose (Life Sciences, TS1M-100). Samples were heated to 72°C for 3 minutes and immediately cooled on ice. Reverse transcription was initiated by adding 6 µL 5× Maxima RT buffer (Thermo Fisher, EP0751), 0.3 µL Maxima RNase H– Minus reverse transcriptase (200 U/µL; Thermo Fisher, EP0751), 10.35 µL 1 M trehalose, 0.3 µL 1 M MgCl₂ (Invitrogen, AM9530G), 0.3 µL of 10 µM template-switching oligo (TSO; 5′-AGACGTGTGCTCTTCCGATCTNNNNNrGrGrG-3′), and 0.75 µL RNase inhibitor (40 U/µL). Reactions were gently mixed and incubated at 50°C for 90 minutes, followed by heat inactivation at 85°C for 5 minutes. For whole-transcriptome amplification, 7 µL of the reverse transcription reaction was added to a PCR master mix consisting of 5 µL nuclease-free water, 0.5 µL 10 µM PCR primer (5′-AGACGTGTGCTCTTCCGATCT-3′), and 12.5 µL KAPA HiFi HotStart ReadyMix (KAPA Biosystems, KK2601). Amplification was carried out for 14 cycles using a 67°C annealing step (20 seconds) and a 6-minute extension at 72°C. Amplified cDNA was purified using a 0.8× AMPure XP bead cleanup (Beckman Coulter, A63881) following the manufacturer’s protocol and eluted in 10 µL elution buffer. DNA concentration was quantified using the QuantiFluor ONE dsDNA System (Promega, E4870) on a Quantus Fluorometer (Promega, E6150), and samples were normalized to 0.2 ng/µL prior to library preparation. Sequencing libraries were constructed using the Nextera XT DNA Library Preparation Kit (Illumina, FC-131-1096), scaled to a final reaction volume of 25 µL. Libraries were normalized by the Heflin Genomics Core via qPCR and sequenced on Illumina NovaSeq 6000 platform.

### Bulk RNA-seq analysis

Reads for 6 dpf *atp1a3a-pt* heads, adult *atp1a3a-pt* brains, and adult *sp4* brains were aligned using the STAR aligner (2.7.3a-GCC 6.4.0-2.28)^109^ to GRCz11 release 104 with the Lawson lab transcriptome annotation version 4.3.2^110^. The number of uniquely mapped reads per sample ranged from 13 to 87 million per adult brain sample, with an average of 41 million. Scripts for differential gene expression analysis using DESeq2^111^ are available on our GitHub: https://github.com/thymelab/BulkRNASeq. For each condition, genes with transcript counts of zero in four or more of the samples across both genotypes were removed, and then raw counts were normalized with rlog counts. Significance was determined with the default DESeq2 method, and both adjusted (Benjamini-Hochberg) and raw *p* values are available in Supplementary Tables S2 to S4.

Downstream RNA-seq analysis was performed using R (Version 4.4.2). Code to reproduce both bulk and single-cell RNA-seq figures is available in Zenodo. Volcano and bar plots were produced using ggplot2 (Version 3.5.2)^112^. Gene ontology (GO) analysis was conducted separately on upregulated and downregulated genes using genes with raw *P* value <0.01 and absolute value of log2 fold change > 0.5 with the “gost” function from gprofiler2 (Version 0.3.2) and FDR correction^113^. Gene set enrichment analysis (GSEA) was performed as previously described^25^ using clusterProfiler (Version 4.14.6)^114^. Gene sets for chromosomal location were created using RefSeq Genes and Gene Predictions from the GRCz11 assembly. For this analysis only, genes were sorted by abs(log2 fold change) prior to GSEA to direct enrichment independent of direction. For all other GSEA, chromosome 19 genes were excluded to minimize artifacts from *cis* effects near the mutant locus, and genes were sorted by log2 fold change. The C5 Molecular Signatures Database^115^ was obtained using the msigdbr package (Version 24.1.0). Gene sets derived from the zebrafish anatomy and development ontology (ZFA) were generated as previously described^25^. For GSEA with terms derived from previously published forebrain single-cell data^49^, markers for all clusters were identified using the “FindAllMarkers” function from Seurat (Version 5.3.0), and the top 250 markers were used to create gene sets. Gene sets for both ZFA and forebrain single cell are available in Zenodo. Network plots were generated using the “emapplot” function from enrichplot (Version 1.26.6)^116^, and heatmaps were generated with the “heatmap.2” function from gplots (Version 3.2.0)^117^. For analysis of previously published forebrain single-cell data^49^, existing clusters were merged to create *snap25a*+ neuronal, radial glia, *ptn*+ astrocyte-like cells, and other clusters. UMAP representations were generated using the “DimPlot” and “FeaturePlot” functions from Seurat (Version 5.3.0). Bubble plot was generated using the “DotPlot” function.

### Data and materials availability

All data are available in the main text, the Supplementary Materials, or appropriate databases. The mutants described in the paper are available from ZIRC. Code is available from https://github.com/thymelab (several repositories, listed throughout the Materials and Methods). Processed behavioral and imaging datasets, additional RNA-seq analysis files and scripts, and juvenile imaging projections are available from Zenodo under DOI 10.5281/zenodo.18672395, and raw behavioral and imaging files that are too large for Zenodo are available upon request. RNA-sequencing data (raw counts and fastq files) is available from GEO under accession number GSE314974.

## Acknowledgments

We thank the Heflin Genomics Institute at UAB, UAB and UMass Chan fish facility staff, and the Research Computing teams at UAB and UMass Chan for supporting this study. This research was funded by the following sources: NIH R00 MH110603 (SBT), Klingenstein-Simons Fellowship Award in Neuroscience (SBT), NARSAD New Investigator Award from the Brain and Behavior Research Foundation (SBT), Pew Biomedical Scholars Award (SBT), and NIH DP2 NS132107 (SBT).

## Author contributions

SBT conceived of the study with input from AJM, analyzed pErk imaging and behavioral data. SBT and AJM wrote the manuscript with contributions from MSC and BLB. AJM analyzed the RNA-seq data. MSC collected all RNA-seq and juvenile brain imaging data, with dissection assistance from JAM. MSC also analyzed the juvenile imaging data. BLB collected the Y-maze juvenile behavioral data, which was analyzed with assistance from CEC. All other authors contributed to the collection of the larval brain imaging and behavioral data, as well as zebrafish line generation and maintenance.

## Notes

### Competing Interest Statement

The authors have declared no competing interest.

