## Supplementary Information for "Zebrafish screen of schizophrenia risk genes reveals convergent dysregulation of cholesterol metabolism"

For Anna J. Moyer *et al.*

**This PDF file includes:**

Supplementary Figs. S1 to S21
Legends for Supplementary Tables S1 to S4

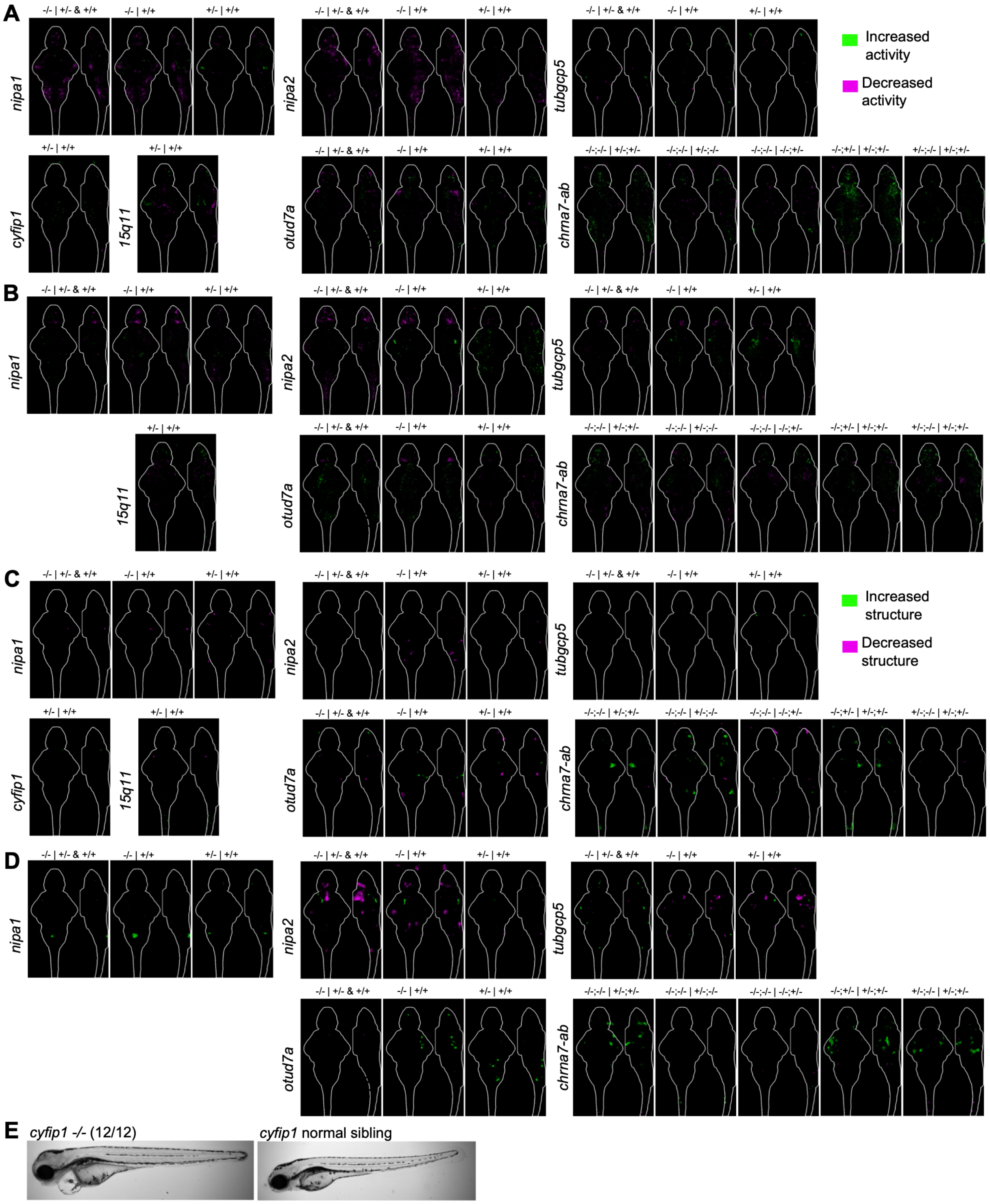


**Fig. S1. Brain activity and structure maps for mutants in genes from CNVs.** Comparisons are shown above the sum-of-slices intensity projection with a 6 dpf brain outline, where the genotype before the | is being compared to the one after. All N are in Table S1. **A**) Brain activity maps. **B**) Replicate set of brain activity maps. **C**) Brain structure maps. **D**) Replicate set of brain structure maps. **E**) Morphological phenotype of *cyfip1* homozygous mutants. The phenotype is shared with the large deletion (15q11).

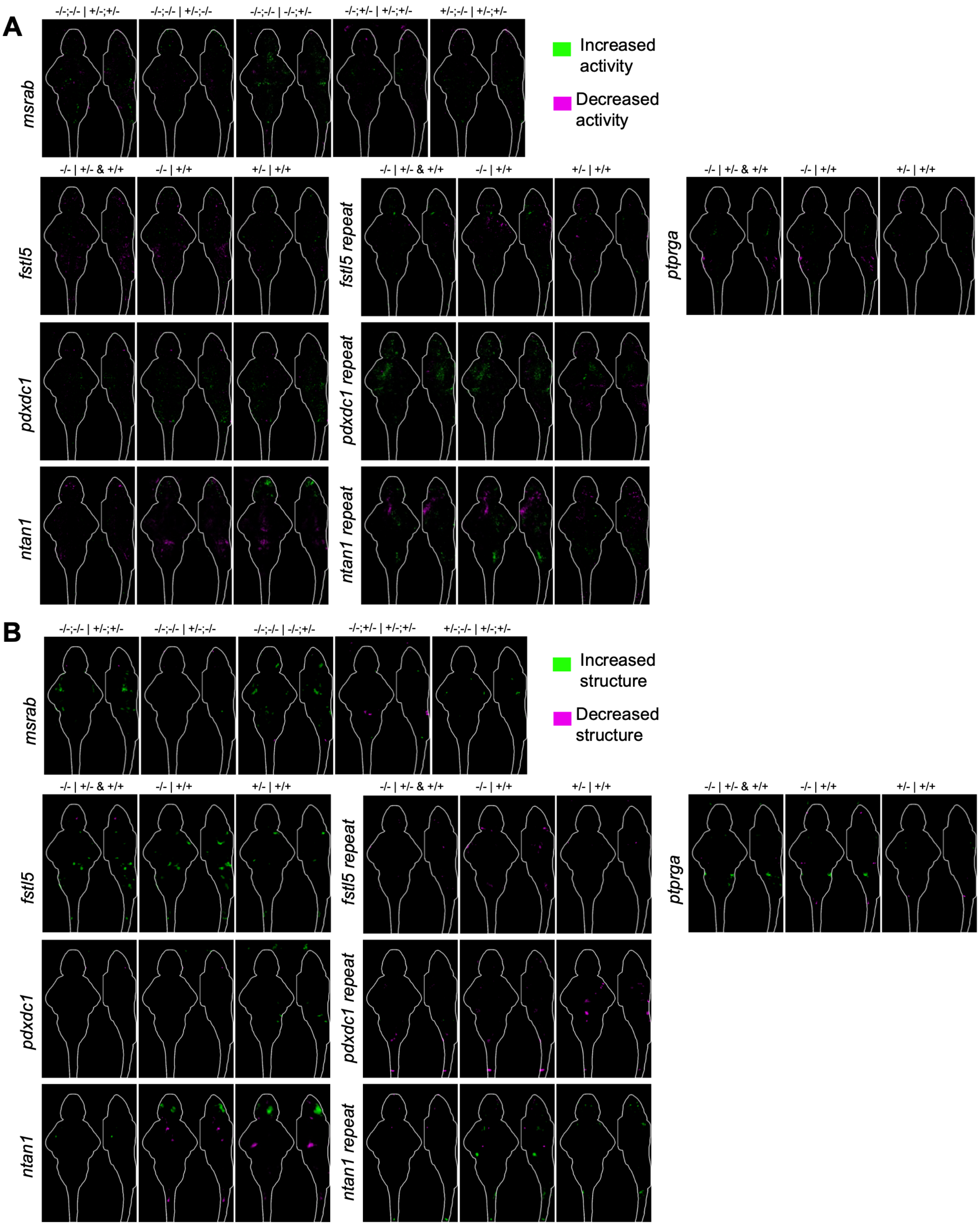


**Fig. S2. Brain activity and structure maps for mutants in rare additional genes/CNVs.** Comparisons are shown above the sum-of-slices intensity projection with a 6 dpf brain outline, where the genotype before the | is being compared to the one after. All N are in Table S1. Repeat experiments are labeled. **A**) Brain activity maps. **B**) Brain structure maps.

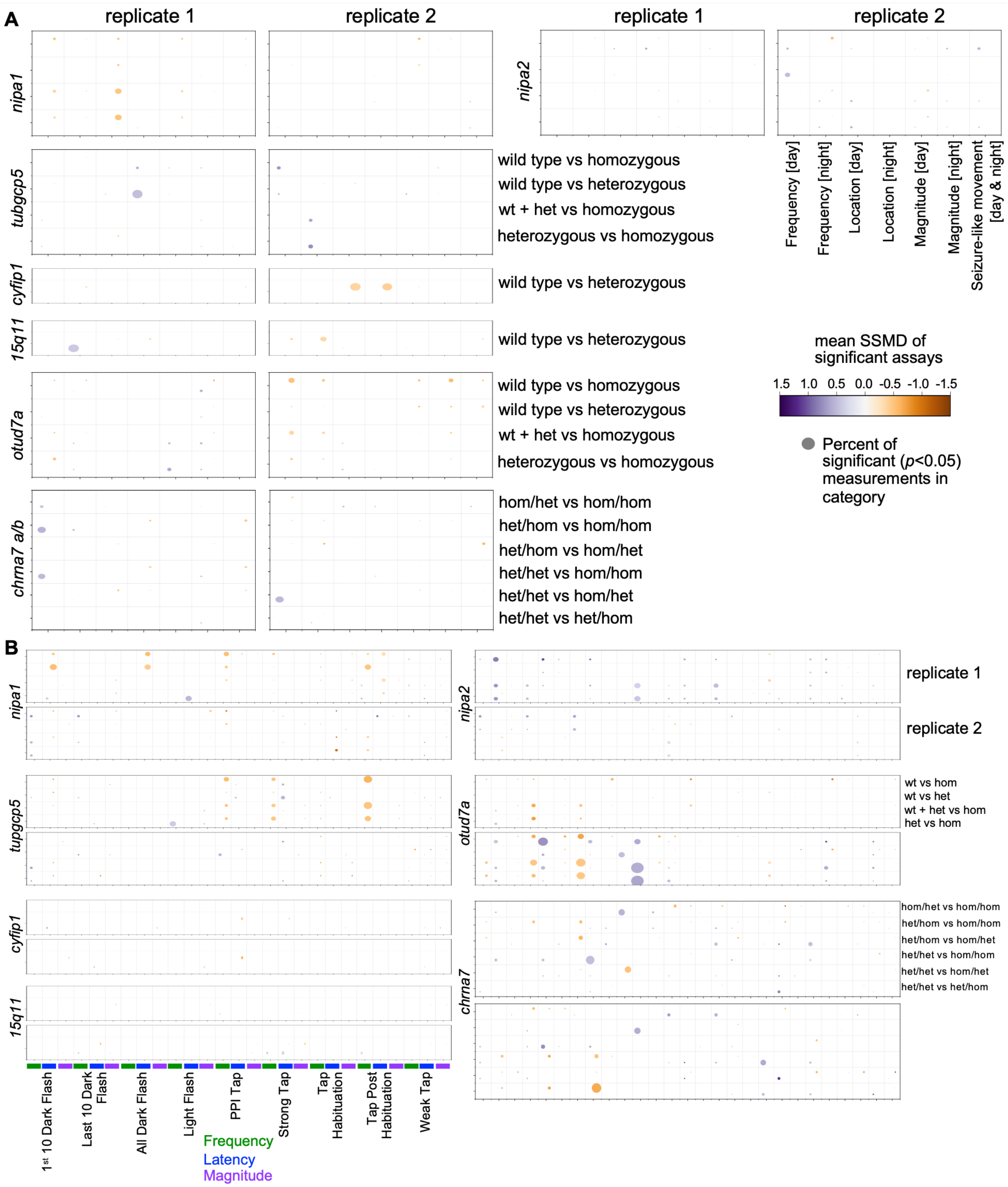


**Fig. S3. Baseline and stimulus-driven behavior summary for mutants in genes from CNVs.** Dot plots for all sibling comparisons. Dot size corresponds to the percent of significant assays in the category (e.g., Magnitude). Replicate experiments using different parental pairs are shown side-by-side. All N are in Table S1. **A**) Baseline behavior. **B**) Stimulus-driven behavior.

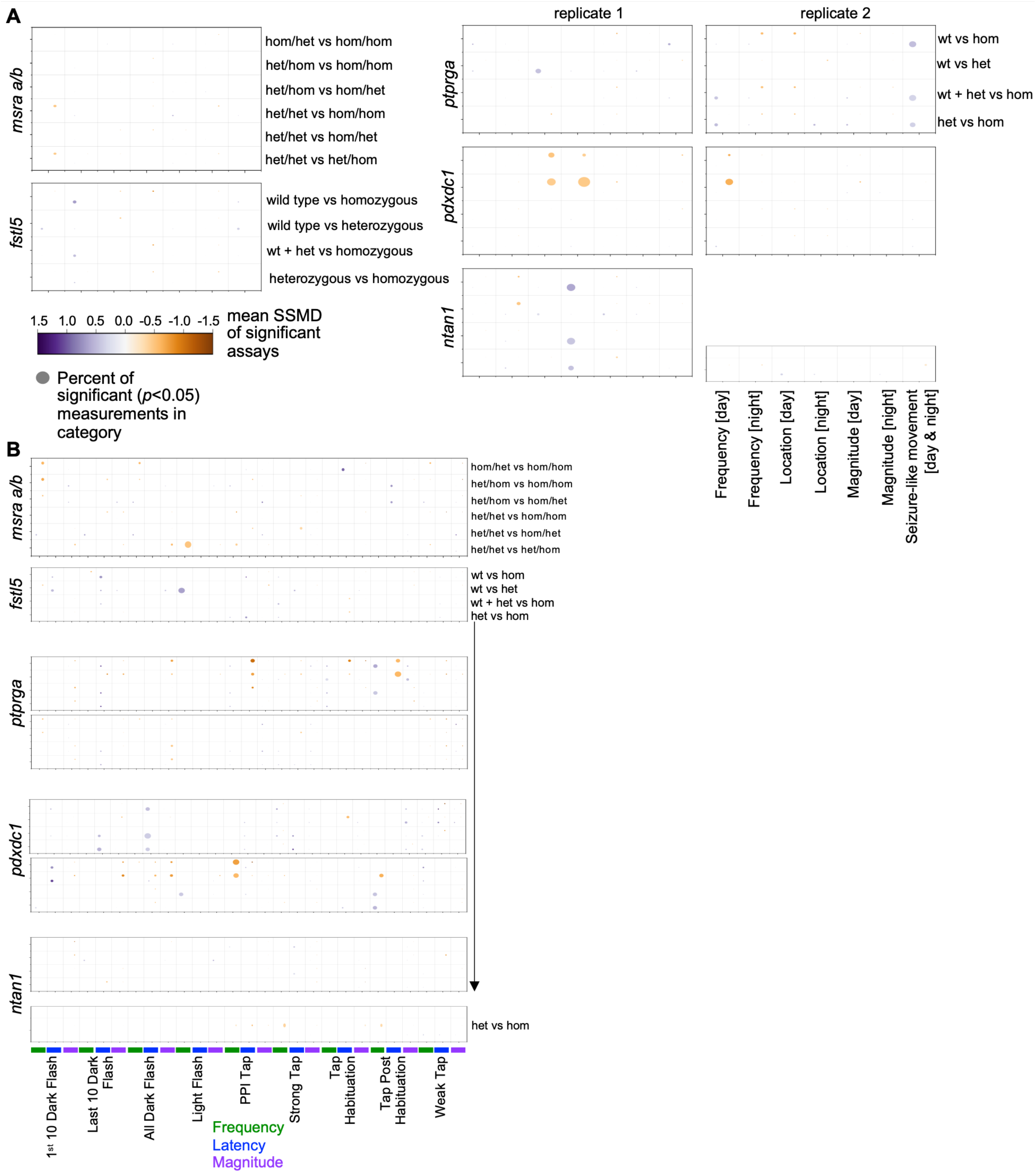


**Fig. S4. Baseline and stimulus-driven behavior summary for mutants in rare additional genes/CNVs.**

Dot plots for all sibling comparisons. Dot size corresponds to the percent of significant assays in the category (e.g., Magnitude). Replicate experiments using different parental pairs are shown side-by-side. All N are in Table S1. **A**) Baseline behavior. **B**) Stimulus-driven behavior.

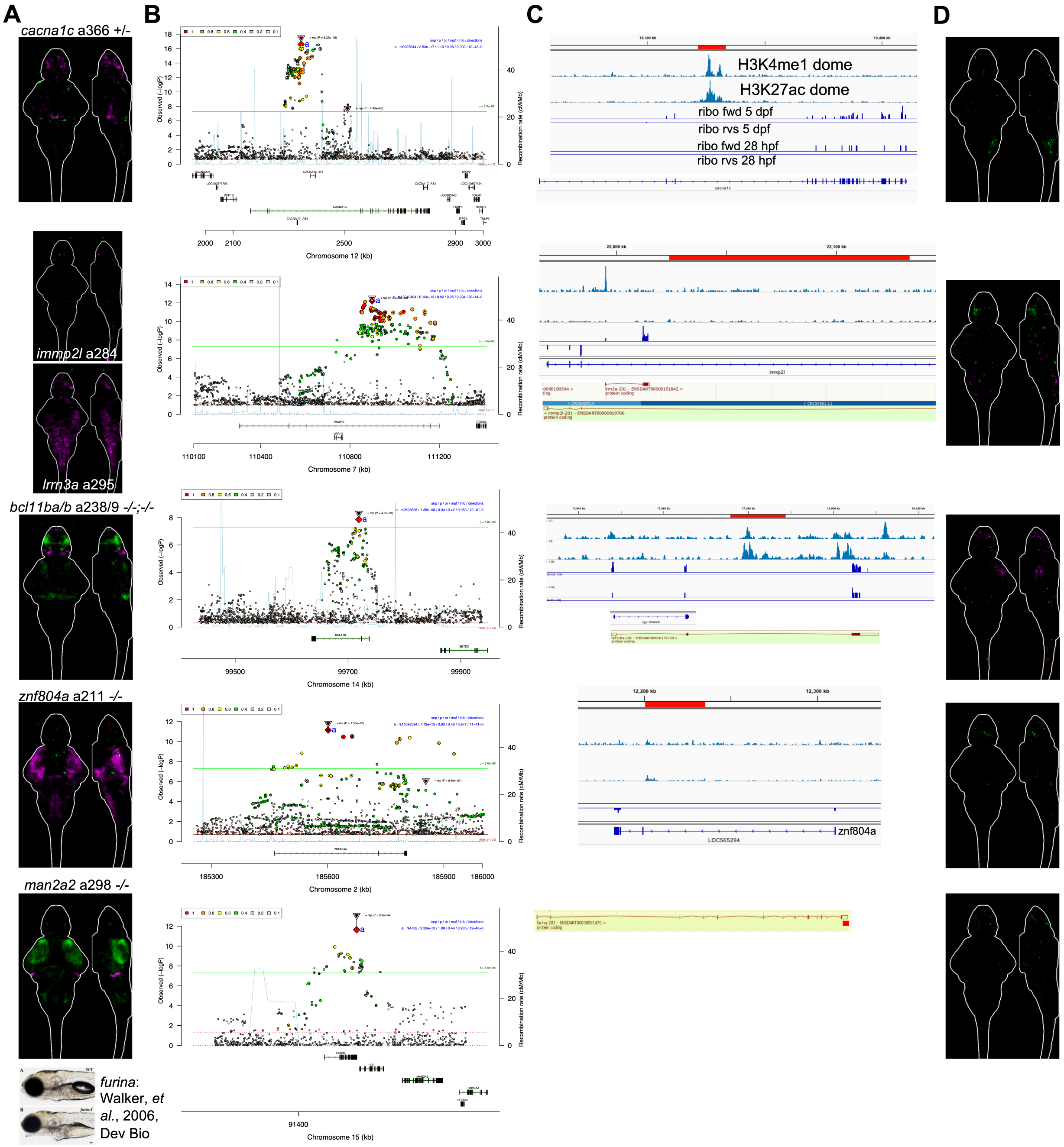


**Fig. S5. Brain activity maps for mutants removing intronic regions.**

Comparisons are shown above the sum-of-slices intensity projection with a 6 dpf brain outline, where the genotype before the | is being compared to the one after. All N are in Table S1. **A**) Original brain activity maps for mutants in the genes from Thyme *et al.*, 2019, except *furina* homozygous mutants, which have a previously published larval morphology phenotype. **B**) Ricopili plot for the GWAS loci. **C**) Visualization of removed region (red). The exact region is available in Table S1. **D**) Brain activity maps for the intronic removal mutants. All are homozygous versus wildtype siblings except for *cacna1c-enh*, which is heterozygous to match the original protein-coding mutation shown in A. The standard behavioral pipeline was also completed and no phenotypes were identified (data not shown, available from Zenodo).

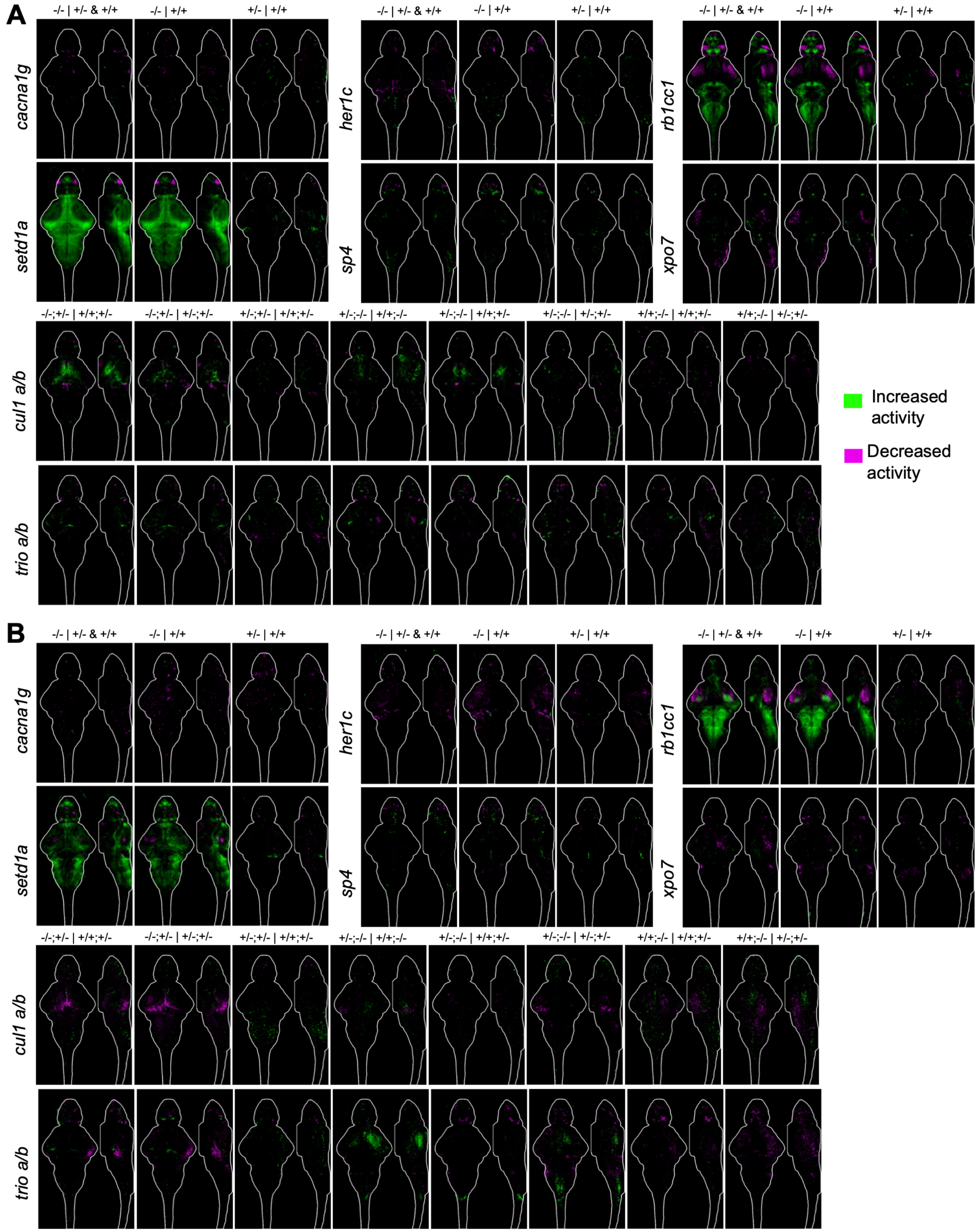


**Fig. S6. Brain activity maps for SCHEMA mutants.**

Comparisons are shown above the sum-of-slices intensity projection with a 6 dpf brain outline, where the genotype before the | is being compared to the one after. All N are in Table S1. Repeat experiments are labeled. **A**) First set. **B**) Replicate set.

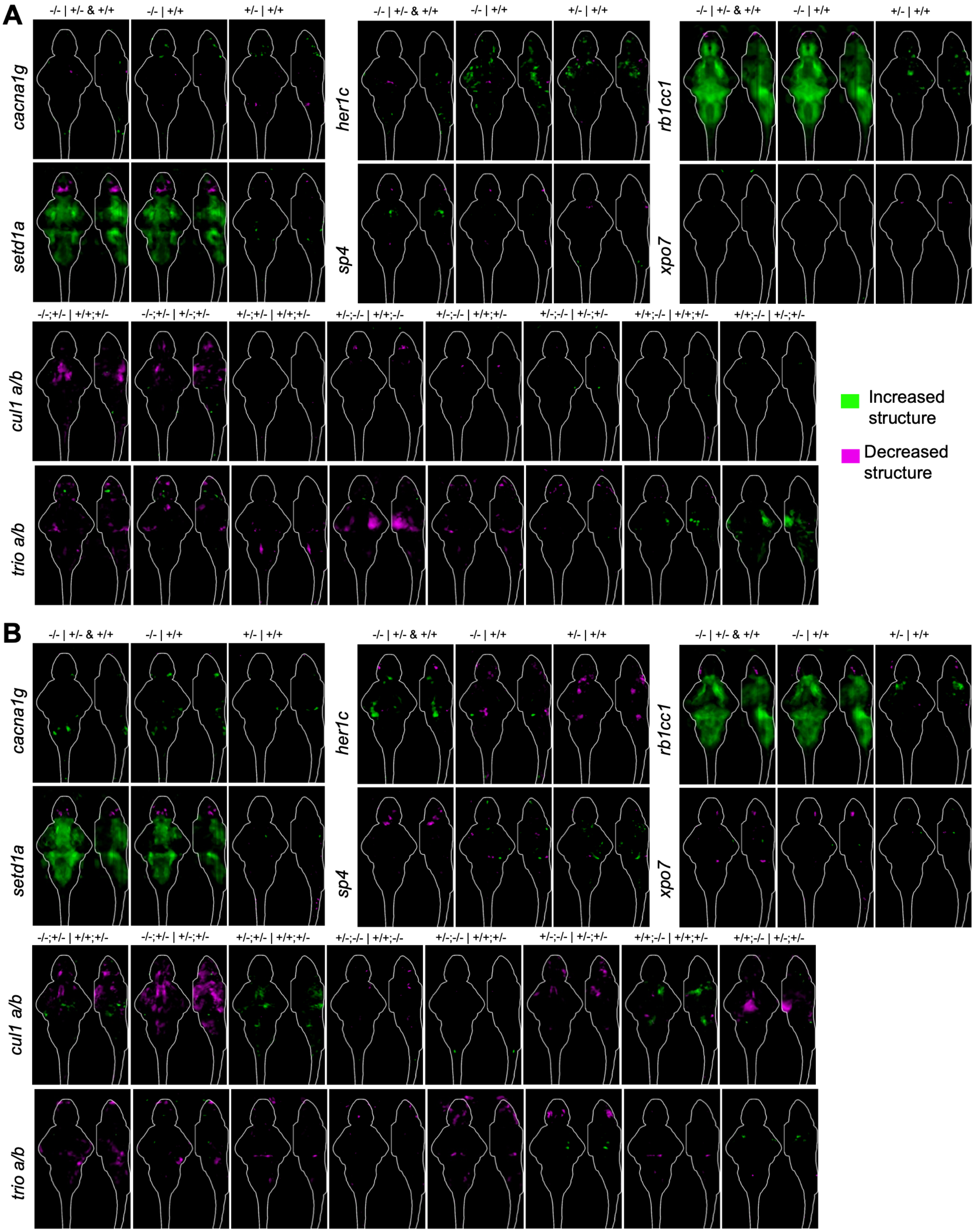


**Fig. S7. Brain structure maps for SCHEMA mutants.**

Comparisons are shown above the sum-of-slices intensity projection with a 6 dpf brain outline, where the genotype before the | is being compared to the one after. All N are in Table S1. Repeat experiments are labeled. **A**) First set. **B**) Replicate set.

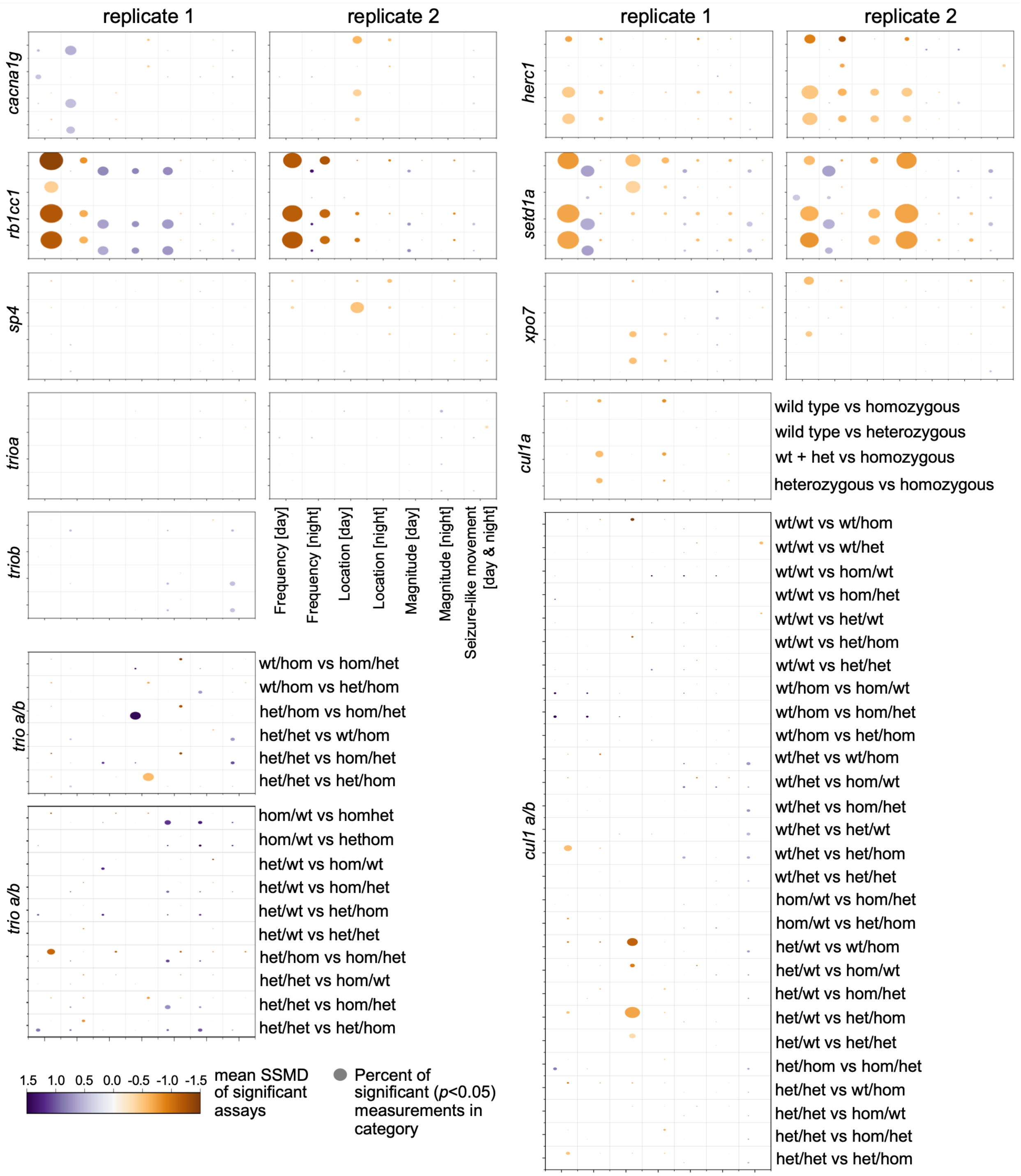


**Fig. S8. Baseline behavior summary data for SCHEMA mutants.**

Dot plots for all sibling comparisons. Dot size corresponds to the percent of significant assays in the category (e.g., Magnitude). Replicate experiments using different parental pairs are shown side-by-side. All N are in Table S1.

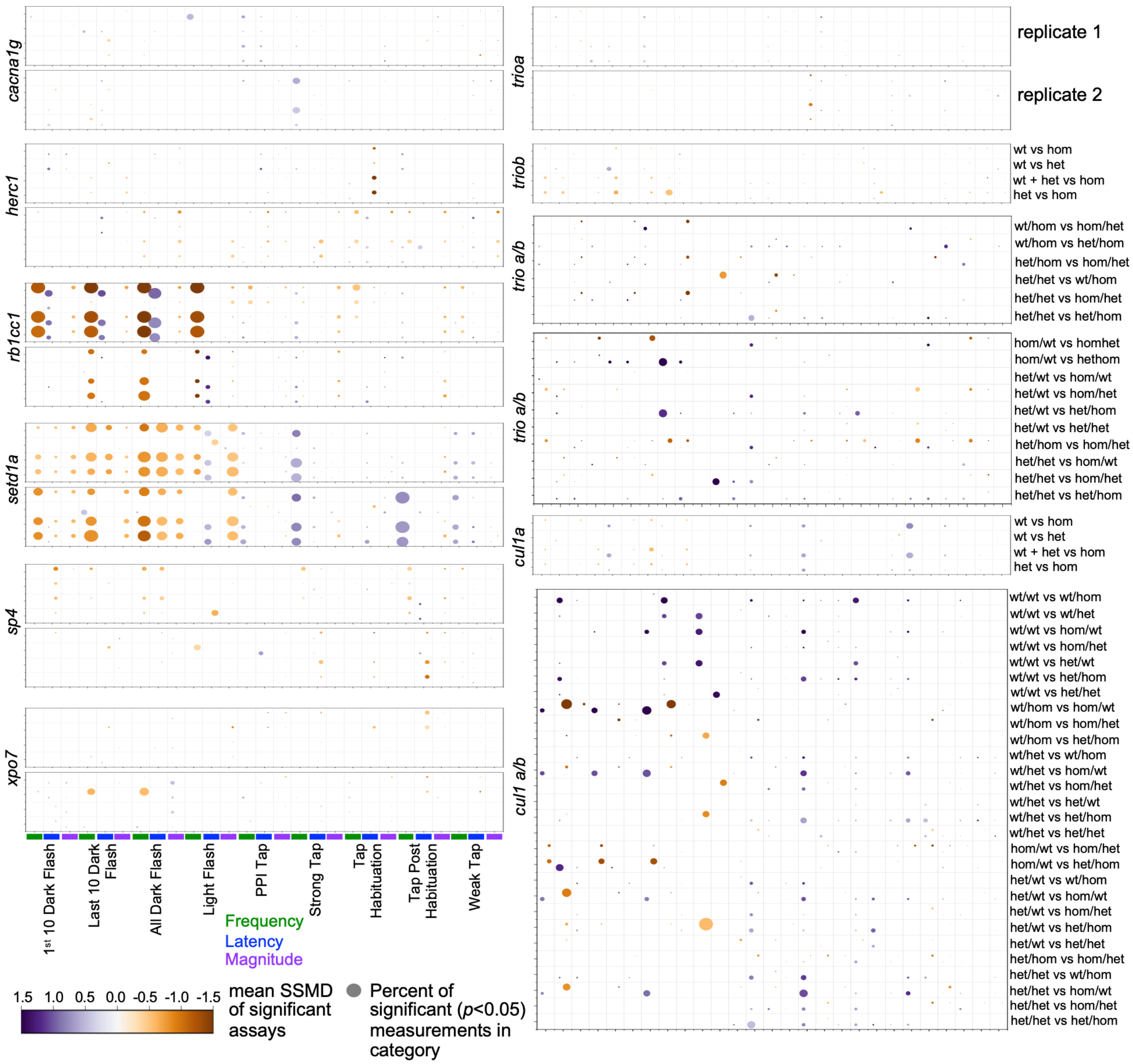


**Fig. S9. Stimulus-driven behavior summary data for SCHEMA mutants.**

Stimulus-driven behavior dot plots for all sibling comparisons. Dot size corresponds to the percent of significant assays in the category (e.g., Weak Tap Magnitude). Replicates are shown top/bottom; N are in Table S1.

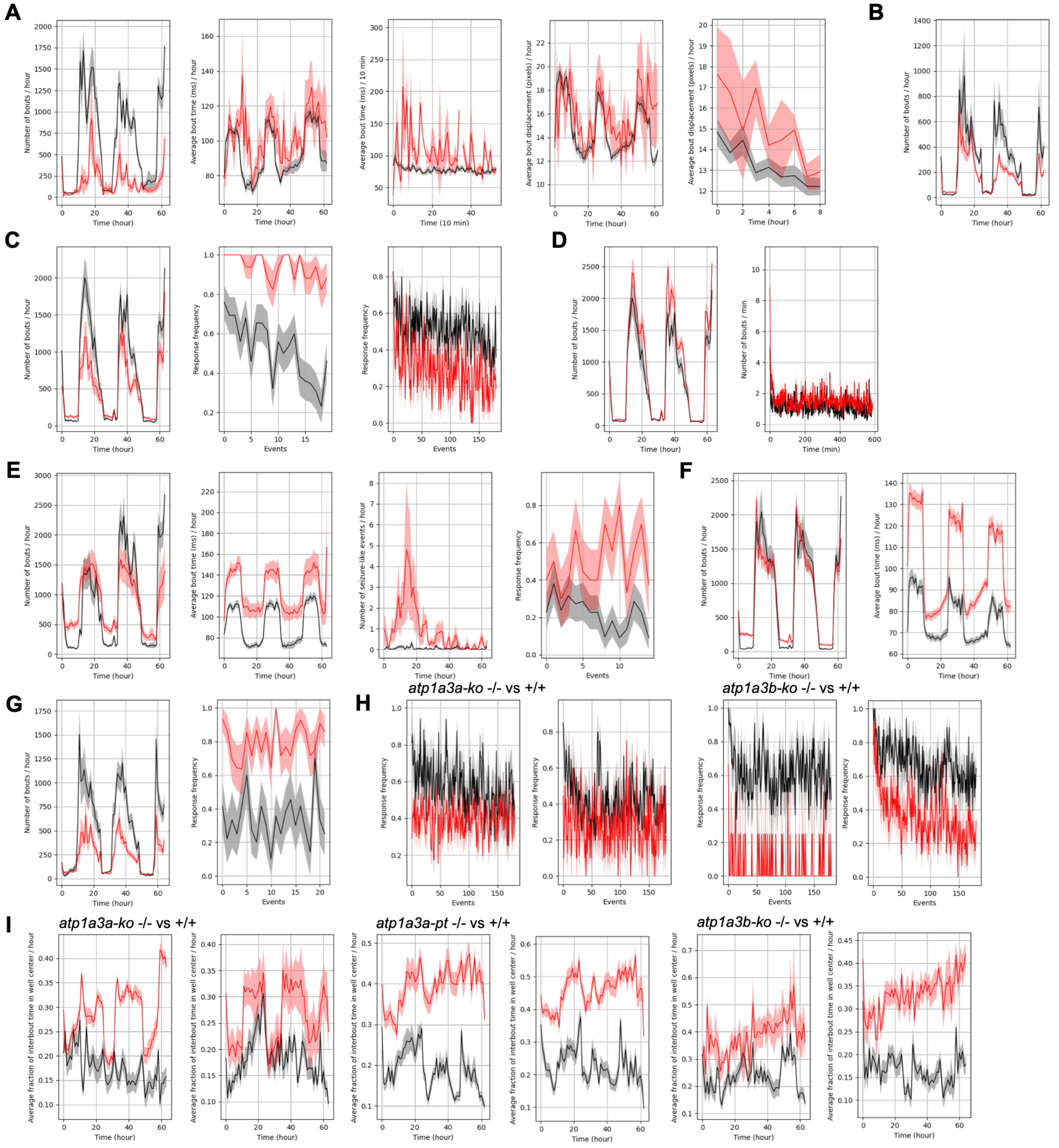


**Fig. S10. Replicate behavioral graphs.**

**A**) Examples of *rb1cc1* baseline behavioral phenotypes. *p* values: 0.00068 (number of bouts / hour), non-significant for bout time for the entire time course and 0.0019 for the day 1 data subsection (day1ppihab, binned per 10-minutes), and non-significant for bout displacement for the entire time course and 0.03 for the day 1 data subsection (day1ppihab, binned per hour). The results are less significant for this set compared to the one in the main text because the N is much lower (see Table S1). **B**) Example of *herc1* baseline frequency of movement phenotype. *p* value: 0.013. **C**) Examples of *setd1a* behavioral phenotypes. *p* values: 0.025 (number of bouts / hour), 1.5e-6 for strong tap response frequency at night at 6 dpf (sound frequency 1400 Hz), 0.0006 dark flash response frequency at 6 dpf (all dark flashes). **D**) No strong phenotype in the *setd1a*

+/- versus +/+ groups, except a weak increased movement frequency on the day 0 night subsection (*p* value: 0.011, binned per minute). **E**) Examples of *atp1a3a-ko* behavioral phenotypes. *p* values: non-significant for entire time course (shown) but significant for most subsections, 0.00018 for bout time, 8.1e-5 for seizure-like movements, and 0.01 for response to light flash response frequency during day of 5 dpf. **F**) Examples of *atp1a3a-pt* behavioral phenotypes. *p* values: 0.015 for bout frequency, 4.1e-5 for bout time. **G**) Examples of *atp1a3b-ko* behavioral phenotypes. *p* values: 5.3e-5 for bout frequency, 2.1e-5 for strong tap response frequency when preceded by a weak prepulse sound (sound frequencies 1400 Hz) during day of 5 dpf. **H**) Dark flash response frequency for all dark flashes for *atp1a3a-ko* and *atp1a3b-ko*. The two biological replicates are shown side-by-side. *p* values: 0.002, 0.035, 0.0053, 4.0e-5. Although graphs are not shown, *p* values for *atp1a3a-pt*: 0.02, 0.01. **I**) Center dwelling preference in the time between bouts for *atp1a3a-ko*, *atp1a3a-pt*, and *atp1a3b-ko* (*atp1a3b-pt* had no phenotype in this measure). The two biological replicates are shown side-by-side. *p* values: 2.3e-7, 0.035, 4.0e-8, 1.1e-8, 0.0037, 3.7e-6.

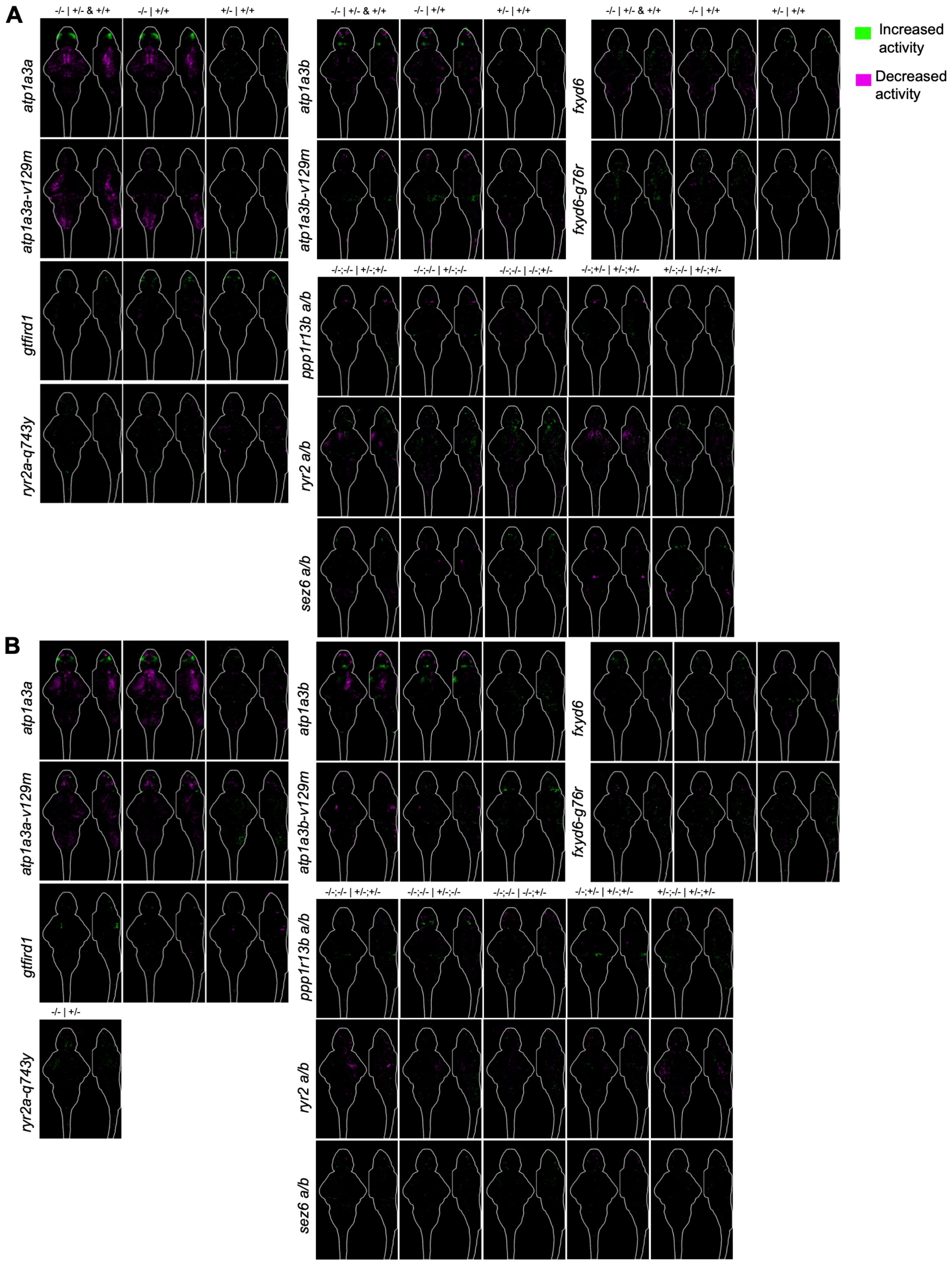


**Fig. S11. Brain activity maps for COS mutants.**

Comparisons are shown above the sum-of-slices intensity projection with a 6 dpf brain outline, where the genotype before the | is being compared to the one after. All N are in Table S1. Repeat experiments are labeled. **A**) First set. **B**) Replicate set.

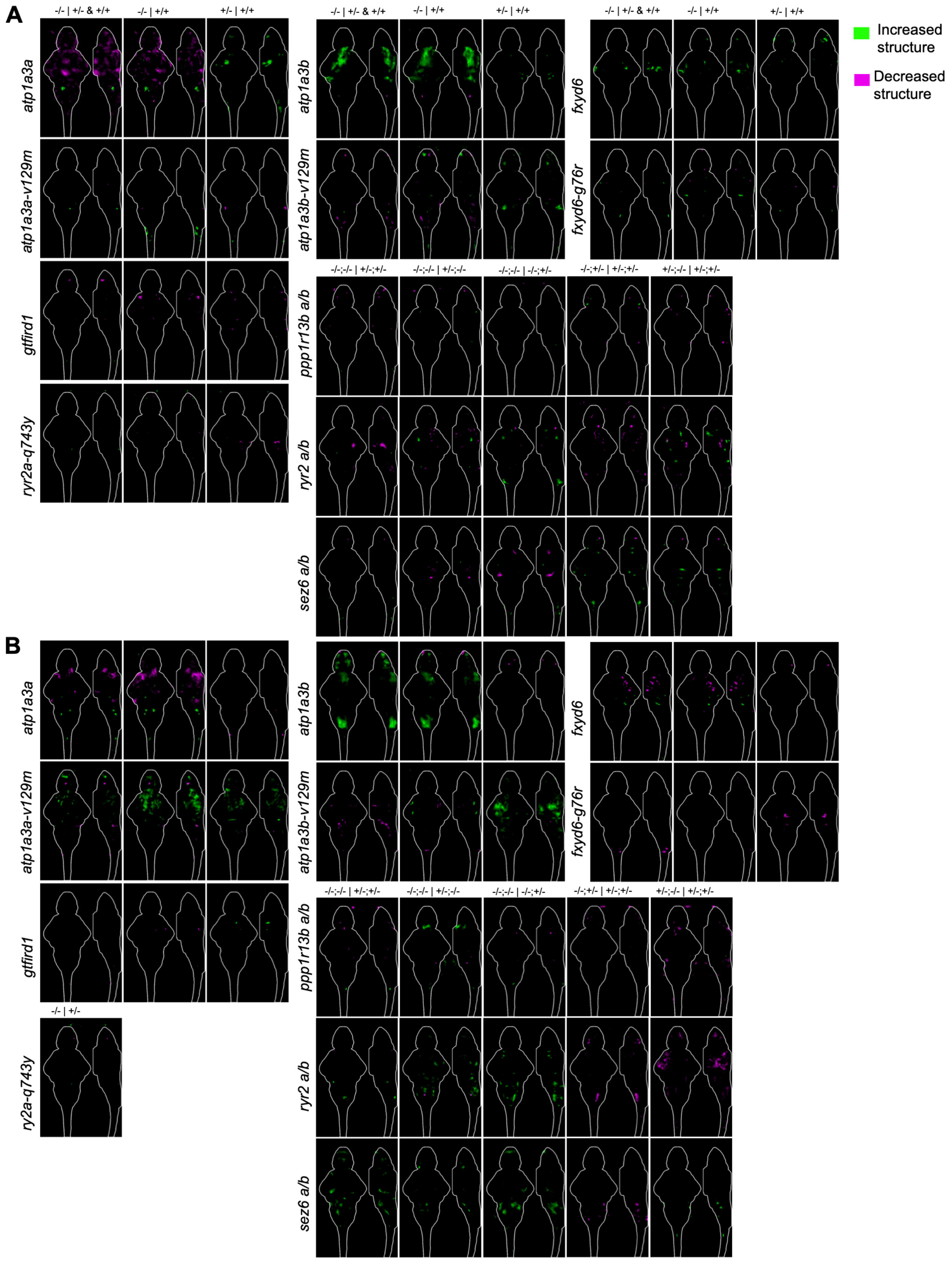


**Fig. S12. Brain structure maps for COS mutants.**

Comparisons are shown above the sum-of-slices intensity projection with a 6 dpf brain outline, where the genotype before the | is being compared to the one after. All N are in Table S1. Repeat experiments are labeled. **A**) First set. **B**) Replicate set.

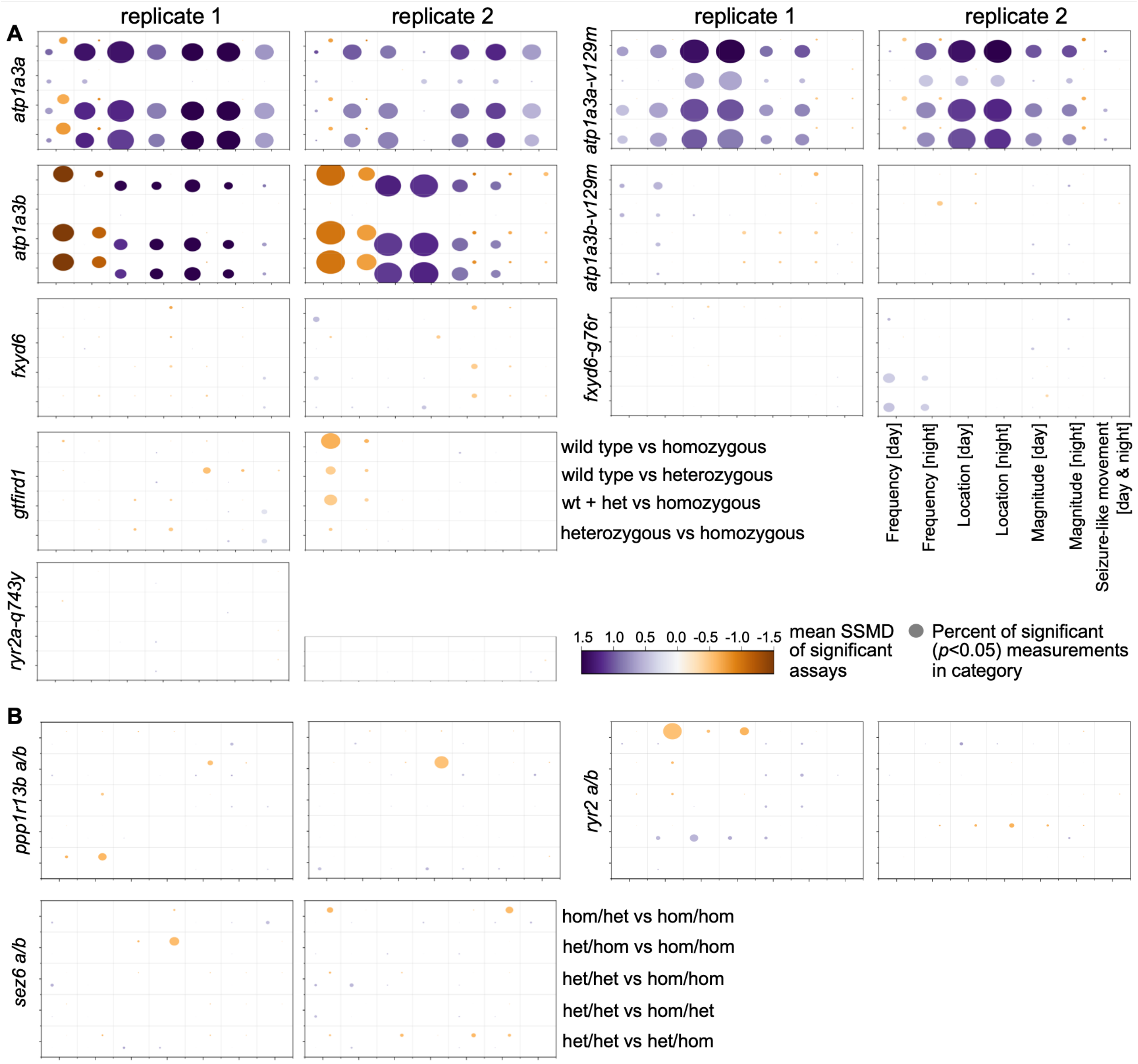


**Fig. S13. Baseline behavior summary data for COS mutants.**

Dot plots for all sibling comparisons. Dot size corresponds to the percent of significant assays in the category (e.g., Magnitude). Replicate experiments using different parental pairs are shown side-by-side. All N are in Table S1.

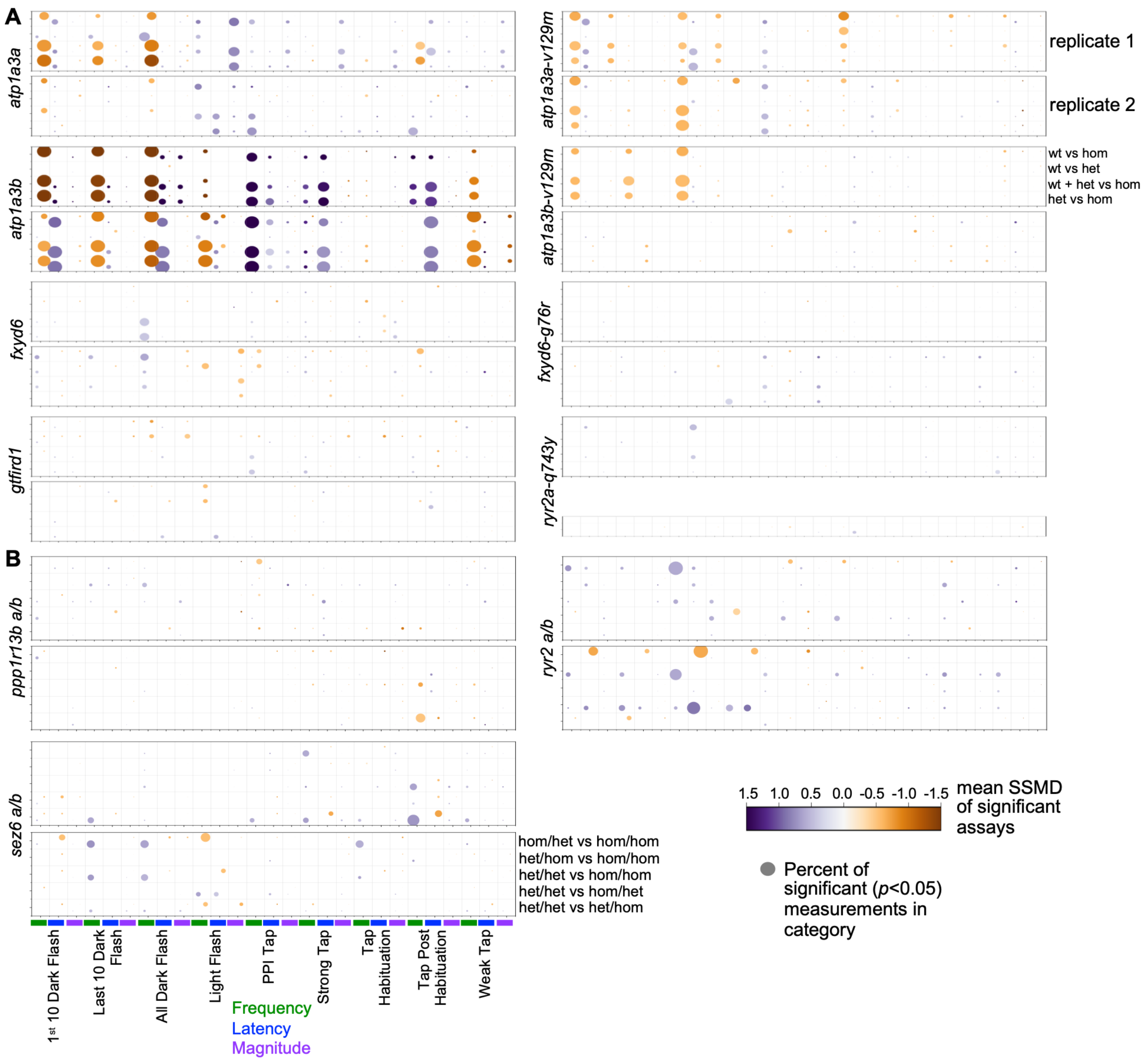


**Fig. S14. Stimulus-driven behavior summary data for COS mutants.**

Stimulus-driven behavior dot plots for all sibling comparisons. Dot size corresponds to the percent of significant assays in the category (e.g., Weak Tap Magnitude). Replicates are shown top/bottom; N are in Table S1.

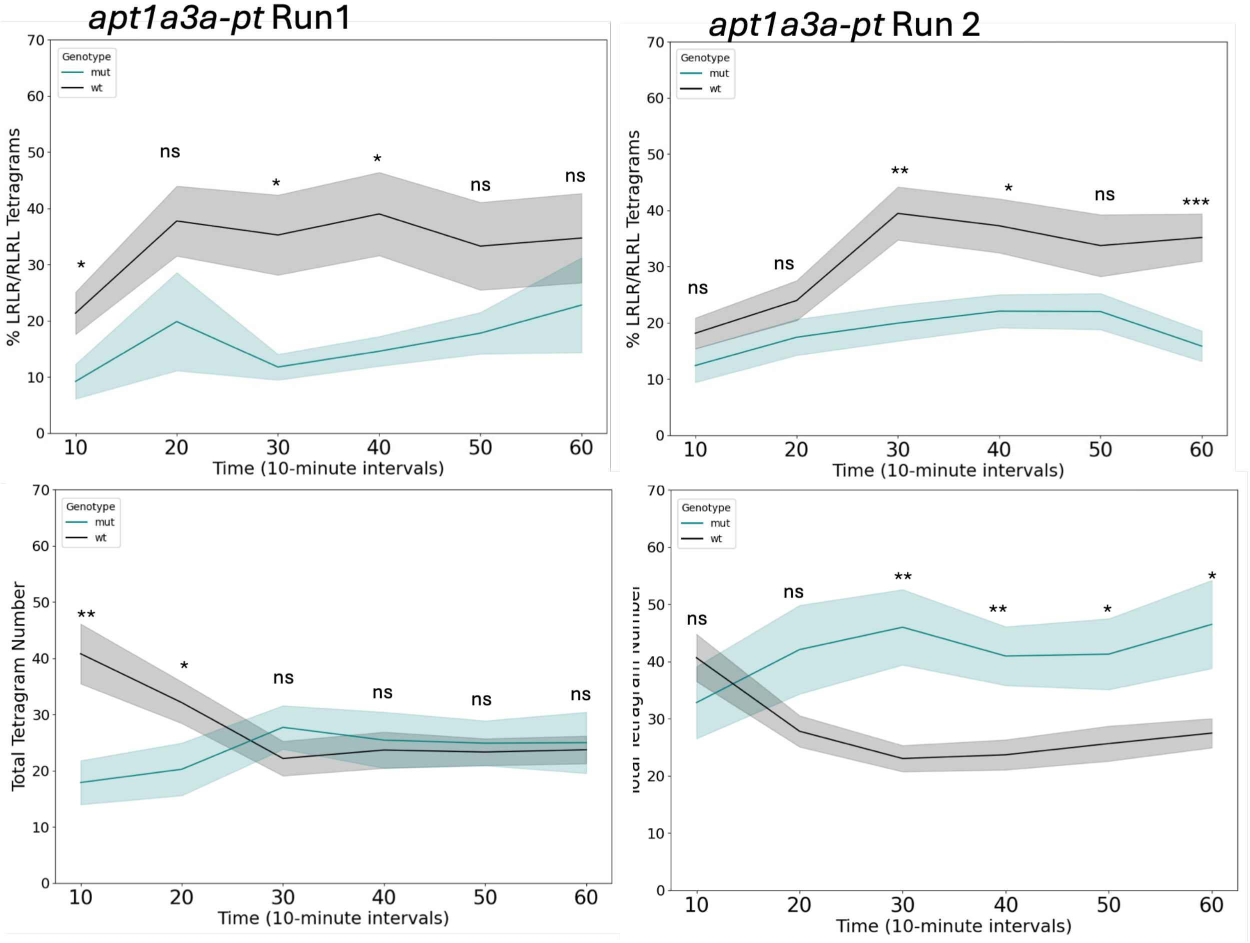


**Fig. S15. Separated Y-maze behavioral assays of *atp1a3a-pt*.**

Percentage of alternation (%LRLR/RLRL) tetragrams (top) and total number of tetragrams performed (bottom) by *atp1a3a-pt* fish in biological replicates 1 and 2 (independent clutches). Means and standard error are plotted at each time point, and each point represents a bin of the previous ten minutes. Significance denotes pairwise comparisons between wildtype (black) and mutant (teal) fish at that individual time point, performed after a Two-Way ANOVA (ns= not significant, * $p$<0.05, ** $p$<0.01, ** $p$<0.001, **** $p$<0.0001). Both individual Two-Way ANOVAs had significant genotype main effects for both alternation percentage tetragrams ($p$<0.001 for each replicate) and total tetragrams ($p$<0.001 for each replicate)

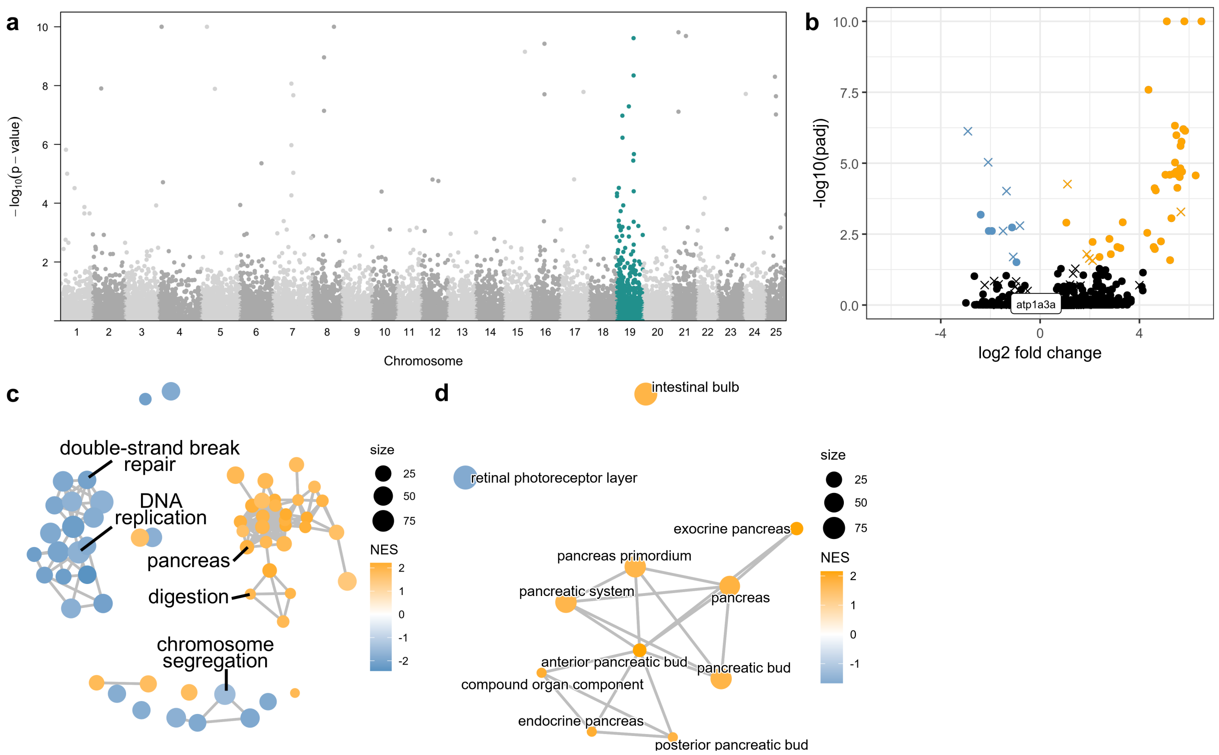


**Fig. S16. Analysis of bulk RNA-seq data from 6 dpf *atp1a3a-pt -/-* heads.**

**A**) Manhattan plot of RNA-seq *p* values from *atp1a3a-pt* homozygous 6 dpf heads plotted in chromosomal order, with GSEA showing no significant enrichment of genes on any chromosome. The *atp1a3a-pt* gene is encoded by chromosome 19 (teal). **B**) Volcano plot of 6 dpf RNA-seq, with upregulated genes labeled in orange and downregulated genes in blue (*padj* <0.05). X indicates transcripts encoded by chromosome 19, and Y-axis values over 10 are plotted at 10. **C**) Network plot with terms identified by C5 GSEA. Nodes represent GSEA terms, and edges represent the number of genes shared between terms. Color indicates the normalized enrichment score, and node size represents term size. **D**) Network plot with terms identified using GSEA derived from the zebrafish anatomy ontology (ZFA).

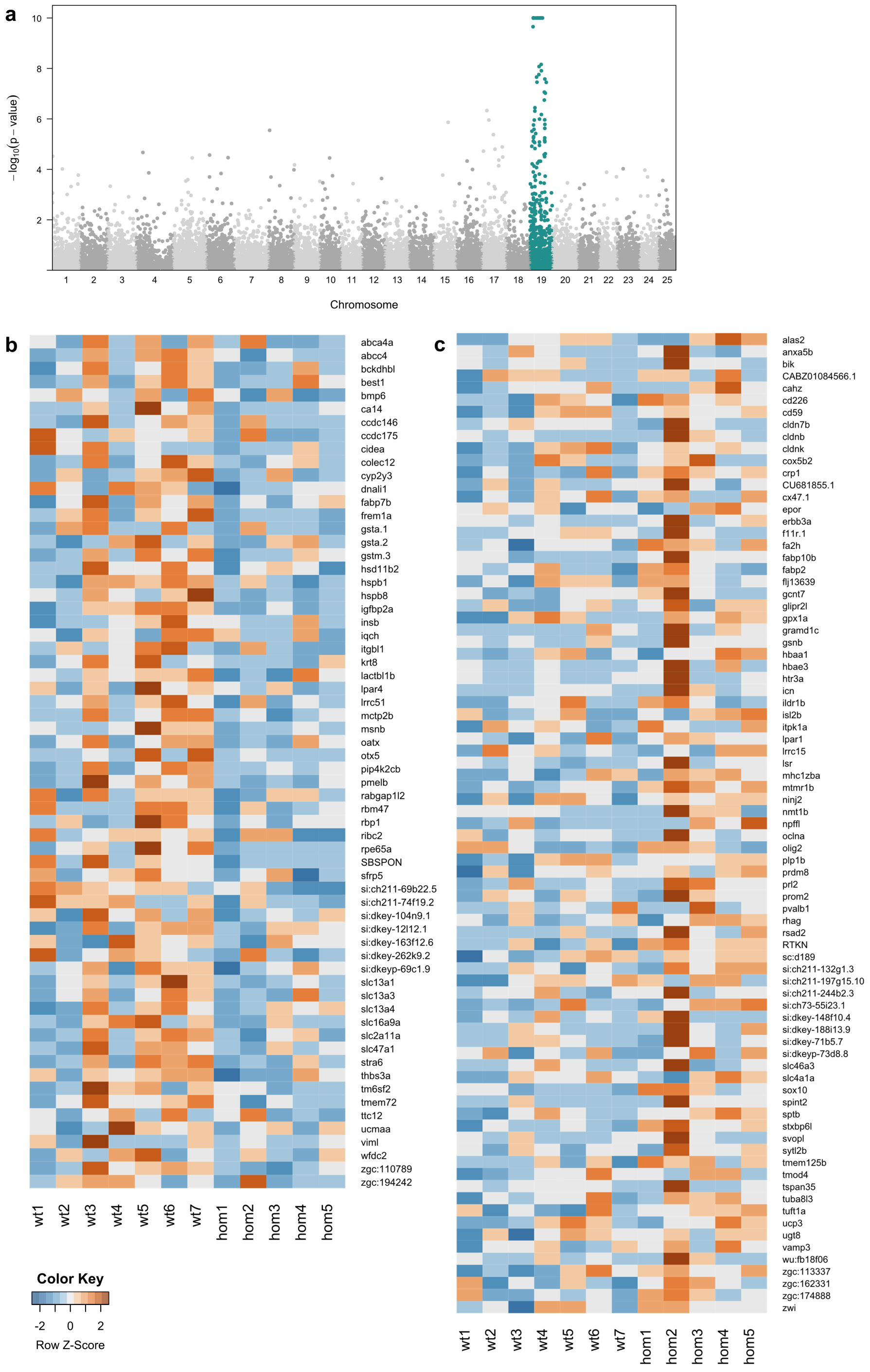


**Fig. S17. Analysis of bulk RNA-seq data from adult *atp1a3a-pt -/-* brains.**

**A**) Manhattan plot of RNA-seq *p* values from *atp1a3a-pt* homozygous adult brains plotted in chromosomal order, showing significant enrichment of chromosome 19, which encodes *atp1a3a-pt* (teal). **B**) Heatmap of genes contributing to negative enrichment of “Progenitor_02 cluster” in GSEA using terms derived from previously published forebrain single-cell data. **C**) Heatmap of genes contributing to positive enrichment of “oligodendrocyte cluster” in GSEA using terms derived from previously published forebrain single-cell data.

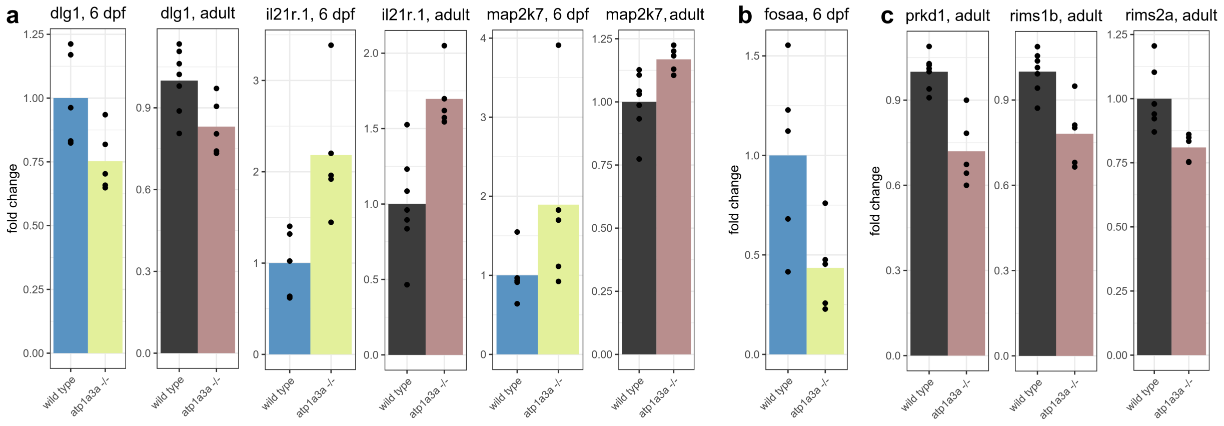


**Fig. S18. Expression of select genes from 6 dpf and adult *atp1a3a-pt -/-* RNA-seq samples.** Bar plots showing fold change for **A**) genes with shared expression between 6 dpf and adult samples, **B**) gene misexpressed only at 6 dpf, and **C**) genes misexpressed only in adult samples. *p* values are in Supplementary Tables S2-3.

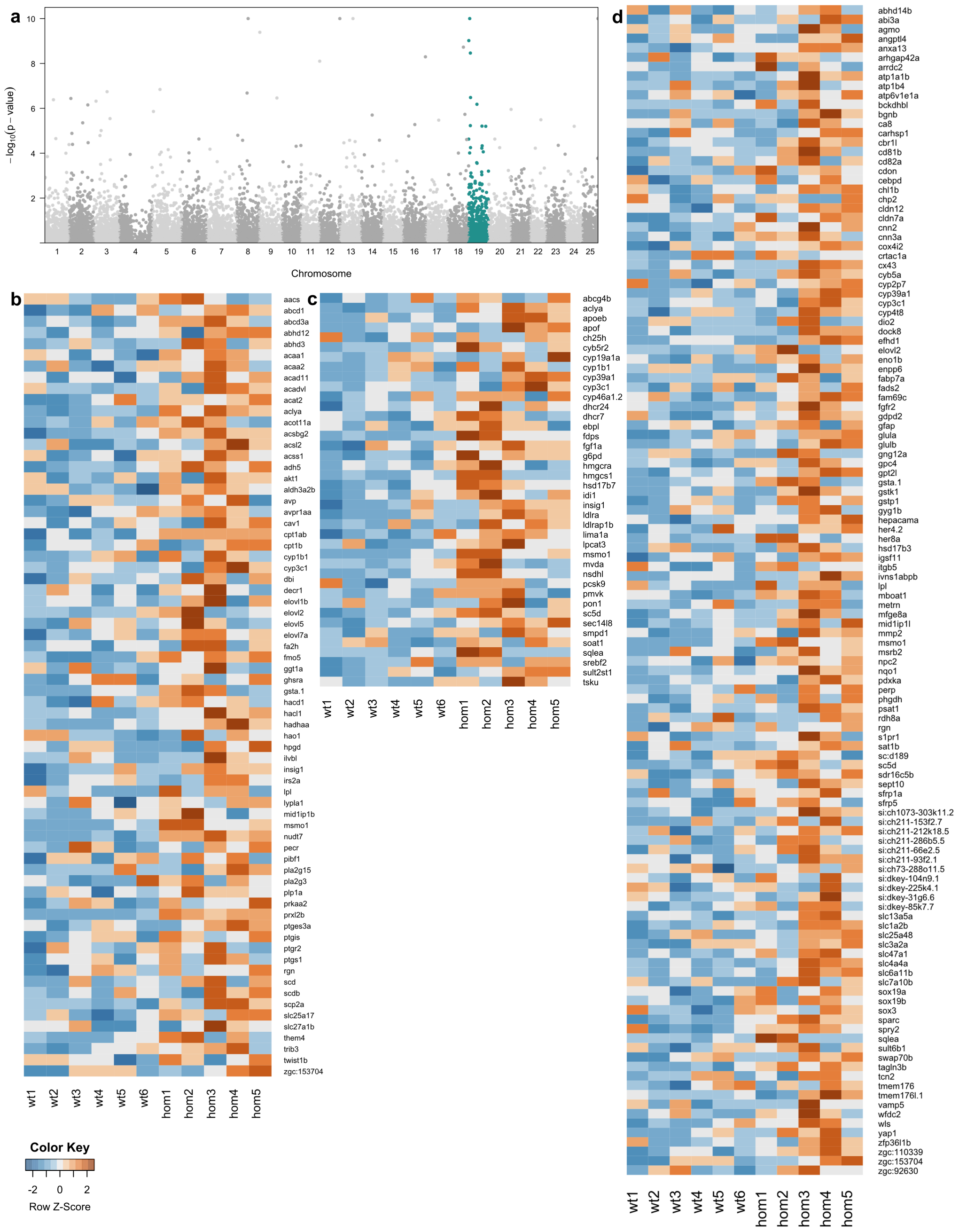


**Fig. S19. Analysis of bulk RNA-seq data from adult *sp4-/-* brains.**

**A**) Manhattan plot of RNA-seq *p* values from *sp4* homozygous adult brains plotted in chromosomal order, showing significant enrichment of chromosome 19, which encodes *sp4* (teal). **B**) Heatmap of genes contributing to positive enrichment of GOBP_FATTY_ACID_METABOLIC_PROCESS term in GSEA using C5 gene sets from the Molecular Signatures Database. **C**) Heatmap of genes contributing to positive enrichment of GOBP_STEROL_METABOLIC_PROCESS term in GSEA using C5 gene sets from the Molecular Signatures Database. **D**) Heatmap of genes contributing to positive enrichment of "astrocyte-like" cluster in GSEA using terms derived from previously published forebrain single-cell data.

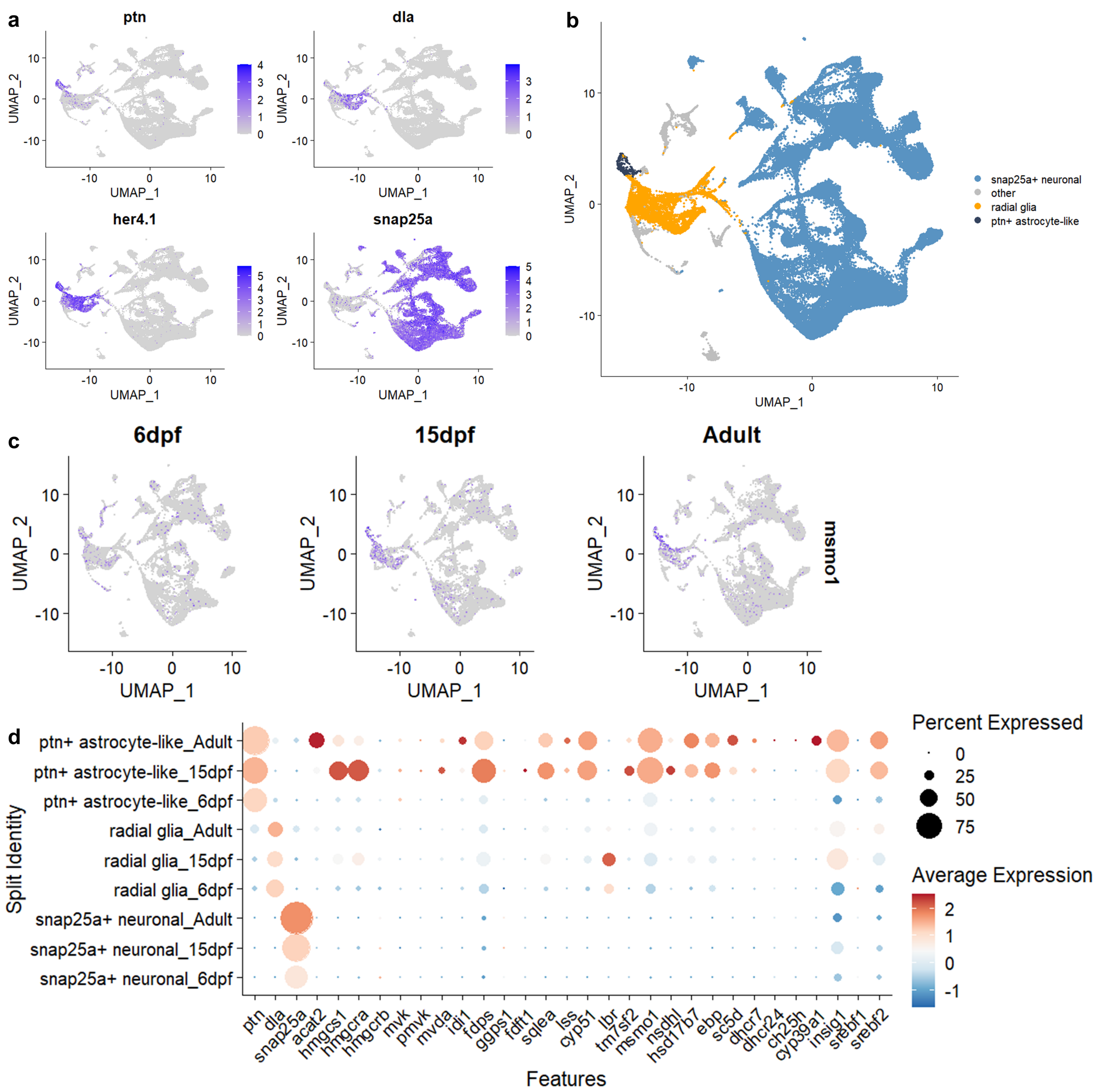


**Fig. S20. Analysis of cholesterol synthesis pathway genes in previously published forebrain single-cell data.**

**A**) UMAP representation of *ptn*, *dla*, *her4.1*, and *snap25a* in forebrain single-cell RNA-seq data. The *ptn* gene is expressed primarily in astrocyte-like cells, *dla* is expressed in radial glia but not in astrocyte-like cells, *her4.1* is expressed in both radial glia and astrocyte-like cells, and *snap25a* is expressed in mature neurons. **B**) UMAP representation of reclustered neural populations to generate a *ptn+/her4+/dla-* astrocyte-like cluster (black), a *ptn-/her4+/dla+* radial glial cluster (orange), and a *snap25a+* neuronal cluster (blue). Non-neural cells (gray) were excluded. **C**) UMAP representations of the cholesterol synthesis gene *msmo1* at 6 dpf, 15 dpf, and adult stages. **D**) Bubble plot showing expression of the marker genes *ptn*, *dla*, and *snap25a* (left) and cholesterol synthesis pathway genes (right) across ages and cell types. Cholesterol synthesis gene expression in astrocyte-like cells appears to increase from 6 dpf to 15 dpf.

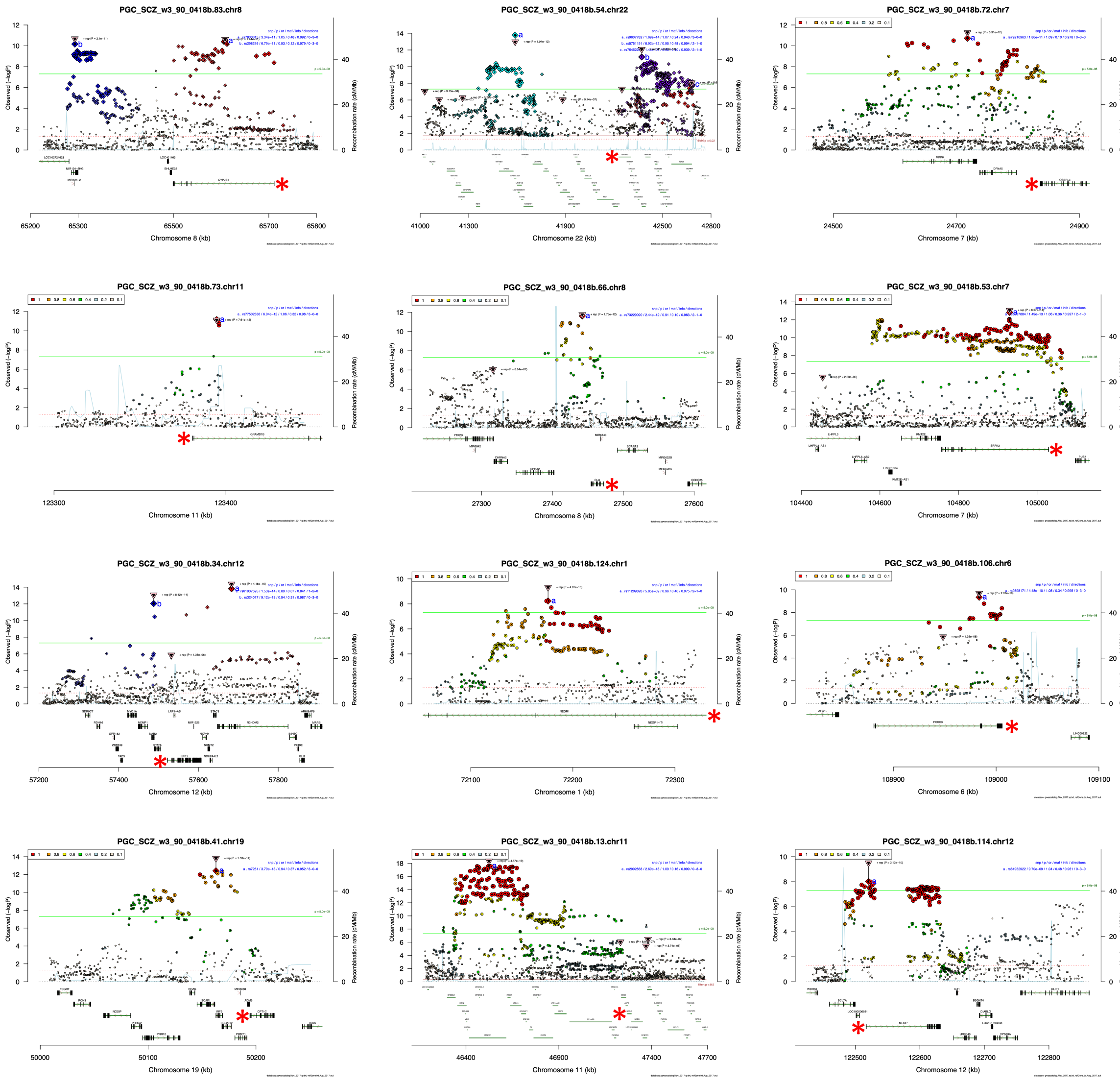


**Fig. S21. Selected RICOPILI plots from the 2022 PGC schizophrenia GWAS.**

These plots include genes with known roles in lipid and cholesterol homeostasis. The gene of interest is marked with a red asterisk.

**Supplementary Table Legends.**

Supplementary Table S1: mutants generated and corresponding genotyping and experimental information.
Supplementary Table S2: DEGs for 6 dpf *atp1a3a-pt* comparing homozygous mutants to wildtype siblings. Other comparisons are available on Zenodo.
Supplementary Table S3: DEGs for adult *atp1a3a-pt* comparing homozygous mutants to wildtype siblings. Other comparisons are available on Zenodo.
Supplementary Table S4: DEGs for adult *sp4* comparing homozygous mutants to wildtype siblings. Other comparisons are available on Zenodo.
